# Staphylococcal protein A inhibits complement activation by interfering with IgG hexamer formation

**DOI:** 10.1101/2020.07.20.212118

**Authors:** Ana Rita Cruz, Maurits A. den Boer, Jürgen Strasser, Frank J. Beurskens, Carla J. C. de Haas, Piet C. Aerts, Guanbo Wang, Rob N. de Jong, Fabio Bagnoli, Jos A. G. van Strijp, Kok P. M. van Kessel, Janine Schuurman, Johannes Preiner, Albert J. R. Heck, Suzan H. M. Rooijakkers

## Abstract

IgG molecules are essential players in the human immune response against bacterial infections. An important effector of IgG-dependent immunity is the induction of complement activation, a reaction that triggers a variety of responses that help to kill bacteria. Antibody-dependent complement activation is promoted by the organization of target-bound IgGs into hexamers that are held together via noncovalent Fc-Fc interactions. Here we show that Staphylococcal protein A (SpA), an important virulence factor and vaccine candidate of *Staphylococcus aureus*, effectively blocks IgG hexamerization and subsequent complement activation. Using native mass spectrometry and high-speed atomic force microscopy, we demonstrate that SpA blocks IgG hexamerization through competitive binding to the Fc-Fc interaction interface on IgG monomers. In concordance, we show that SpA interferes with the formation of (IgG)_6_:C1q complexes and prevents downstream complement activation on the surface of *S. aureus.* Lastly, we demonstrate that IgG3 antibodies against *S. aureus* can potently induce complement activation even in the presence of SpA. Altogether, this study identifies SpA as an immune evasion protein that specifically blocks IgG hexamerization.

## Introduction

Antibodies play a key role in the human immune response against bacterial infections. While antibodies can bind and neutralize bacterial virulence factors, they can also signal to components of the innate immune system and induce bacterial killing. To do so, antibodies bind bacterial cells via their variable (Fab) region and subsequently trigger Fc-mediated effector functions (Lu *et al*, 2018). The complement system, a large network of plasma proteins, forms an important effector of antibody-dependent immune protection against invading bacteria. An activated complement cascade results in efficient decoration of bacteria with C3-derived molecules that are essential to trigger highly effective phagocytic uptake via complement receptors on phagocytes. Furthermore, complement generates chemoattractants and induces direct killing of Gram-negative bacteria. Because effective complement activation is an important effector mechanism of therapeutic antibodies in cancer (Lee et al, 2017), the ability of complement to kill bacteria could also be exploited for antibacterial therapies against (antibiotic-resistant) pathogens (Laxminarayan *et al*, 2013; Theuretzbacher & Piddock, 2019; Sause *et al*, 2016).

The antibody-driven, ‘classical’ complement pathway is initiated when circulating C1 complexes are recruited to antibody-labeled target surfaces (Gaboriaud *et al*, 2004). The most abundant antibody isotype in serum is IgG, which is subdivided into subclasses IgG1, IgG2, IgG3 and IgG4, in order of decreasing abundance. IgG antibodies can bind surface antigens via their Fab regions and subsequently recruit C1 via their Fc region (Fig EV1A). The C1 complex consists of three large units: C1q, C1r and C1s. C1q comprises the antibody recognition unit of the C1 complex and is composed of six globular heads connected by collagen-like stalks (Strang *et al*, 1982). Upon binding of C1q, its associated proteases C1r and C1s are activated to cleave other complement proteins that together form enzymes on the surface that catalyze the covalent deposition of C3b molecules onto the bacterial surface (Fig EV1A). C3b molecules are recognized by complement receptors on phagocytes (neutrophils, macrophages), which engulf and digest bacteria intracellularly. The deposition of C3b also results in amplification of the complement cascade and the activation of downstream complement effector functions.

In the past years, it has become clear that efficient binding of C1 to target-bound IgG molecules requires IgGs to form ordered hexameric ring structures (Diebolder *et al*, 2014; Strasser *et al*, 2019). Cryo-electron tomography and atomic force microscopy studies revealed that the six globular heads of C1q can simultaneously bind to each of the six IgG molecules that form a hexameric binding platform (Diebolder *et al*, 2014) (Fig EV1B). The formation of these hexamers is induced upon antibody binding to surface-bound antigens and driven by non-covalent interactions between the Fc regions of neighboring IgG molecules (Strasser *et al*, 2020) (Fig 1A).

**Figure 1.**
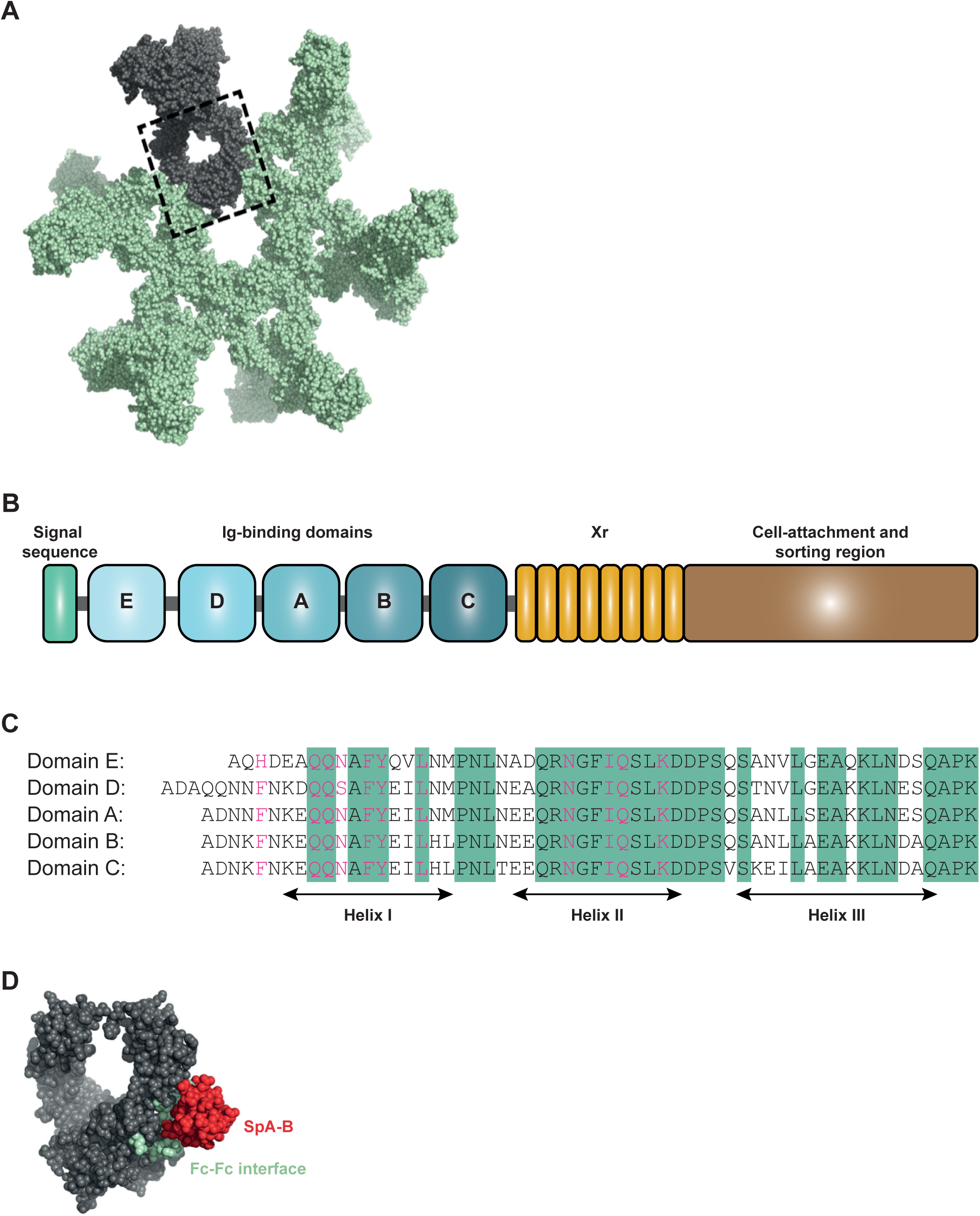
The Ig-binding domains of staphylococcal protein A (SpA) bind to residues of the IgG-Fc region that are involved in IgG hexamerization. A IgG hexamer crystal packing of IgG1-b12 (pdb 1HZH). A single IgG is depicted in grey and IgG-Fc domain is enclosed in the dashed box. B Schematic representation of SpA organization. SpA consists of a signal sequence, five immunoglobulin (Ig)-binding domains (E, D, A, B, and C), an Xr region (octapeptide repeats variable in number), and a cell-wall attachment and sorting region that includes a constant Xc region, the LPETG motif, a hydrophobic anchor and positively charged residues. C Sequence alignment of the five highly homologous Ig-binding domains of SpA. The amino acid residues conserved in all five domains are highlighted in green. The residues involved in the interaction with the Fc region of IgG are colored in pink. D Space-filling presentation depicting the Fc domain of IgG1-b12 and its interaction with the B-domain of SpA (SpA-B: pdb 1FC2; complementary Fc docking domain for SpA-B is hidden). The residues involved in Fc-Fc interactions required to form the IgG hexameric ring are depicted in green and the crystal structure of SpA-B is shown in red.

Interestingly, some bacteria produce IgG-binding molecules that recognize the Fc-domain of IgGs (Sidorin & Solov’Eva, 2011). Best-known is Staphylococcal protein A (SpA), a 42 kDa protein that has a high affinity for the Fc region of IgG and that is therefore commonly used as a tool in affinity-chromatography to purify monoclonal antibodies (Hober *et al*, 2007). SpA is produced by *Staphylococcus aureus* (*S. aureus*) (Forsgren, 1970; Peacock *et al*, 2002), an important human pathogen which is the main cause of serious hospital-acquired infections such as bacteremia, sepsis and endocarditis (Lowy, 1998). Due to the dramatic increase of antibiotic resistance and the lack of proper vaccines, physicians are frequently left with no useful or suboptimal alternatives when treating these infections.

SpA is considered an important virulence factor (Kim *et al*, 2011; Palmqvist *et al*, 2002; Gómez *et al*, 2004; Hong *et al*, 2016; Patel *et al*, 1987) and vaccine candidate (Falugi *et al*, 2013; Kim *et al*, 2010). The protein is abundantly present on the bacterial cell wall (Gatlin *et al*, 2006; Ventura *et al*, 2010; Ythier *et al*, 2012) but also released in the extracellular environment (Becker *et al*, 2014; O’Halloran *et al*, 2015). SpA is composed of a signal sequence, five sequential Ig-binding domains (denoted E, D, A, B, and C), an Xr region and a cell-wall attachment and sorting region (Uhlen *et al*, 1984) (Fig 1B). Each of the five repeating Ig-binding domains adopts a three-helical structure that can bind to the Fc region of IgG via helices I and II (Deisenhofer, 1981) and the Fab region of VH3-type family of antibodies via helices II and III (Sasso *et al*, 1989, 1991; Graille *et al*, 2000). Binding of SpA to Ig-Fc regions is described to protect *S. aureus* from phagocytic killing (Falugi *et al*, 2013) while cross-linking of Ig-Fab regions triggers proliferation and apoptotic collapse of B cells (Goodyear & Silverman, 2003). The five Ig-binding domains are highly homologous, sharing 74 to 91% of their amino acid sequence (relative to the A domain) (Deis *et al*, 2014) (Fig 1C). It was demonstrated that the binding interface of the B domain of SpA (SpA-B) and IgG1-Fc involves 11 amino acid residues of SpA-B and 9 residues of IgG1-Fc (Deisenhofer, 1981). Interestingly, SpA binds to all IgG subclasses except for IgG3, due to a substitution in one of the nine Fc-contact residues in IgG3 (His^435^ in IgG1 becomes Arg^435^ in IgG3) (Jendeberg *et al*, 1997). The reported crystal structure of SpA-B and its IgG-Fc interaction site are depicted in Fig 1D. Noteworthy, SpA binds to the same interface where the IgG Fc-Fc interactions take place to form the hexameric IgG platform required for complement activation.

Here, we investigate the impact of IgG-Fc binding properties of SpA on the assembly of IgG molecules into hexamers, both in solution and on antigenic surfaces. We show that SpA blocks the formation of the hexameric C1q binding platform and as a result inhibits IgG-dependent complement activation. Our data provide an important contribution to the understanding of molecular mechanisms of complement evasion, which is crucial for the intelligent design of new therapeutic strategies to tackle infectious diseases.

## Results

### Staphylococcal protein A (SpA) binds to the Fc region of IgG1, IgG2 and IgG4 with 1:1 stoichiometry

To investigate whether the interaction between SpA and IgG-Fc could affect IgG-dependent complement activation, we first examined the IgG-binding properties of SpA. Most *S. aureus* strains express SpA with five highly homologous Ig-binding domains (A-E) (Fig 1C) that each can bind to the Fc region of IgG. However, the single B-domain of SpA has been extensively used for structural and biochemical studies (Deisenhofer, 1981; Gouda *et al*, 1998). Thus, here we tested both a soluble SpA construct containing all five domains (SpA) and a SpA construct consisting solely of the B-domain (SpA-B). To exclude potential interactions between the SpA proteins and IgG Fab domains, we used human monoclonal IgGs that do not bind SpA via the Fab region. Accordingly, SpA-B^KK^ (SpA-B domain with abolished Fc-binding, but intact Fab-binding properties (Falugi *et al*, 2013)) was found not to interact with the IgGs used in this study (Fig EV2).

We used ELISA and native mass spectrometry (native MS) to study the interaction between the SpA constructs and human IgGs. Native MS analyzes masses of intact protein complexes in a native state, allowing non-covalent interactions to remain intact (Leney & Heck, 2017; Wang *et al*, 2016). In line with previous studies, our results indicated that both SpA constructs bind strongly to the Fc region of all human IgG subclasses, except IgG3 (Fig 2 and EV3, see Table EV1 for the determined molecular weights (MW) of all proteins analyzed by native MS). Interestingly, the binding stoichiometry differed between SpA and SpA-B. As expected with the presence of two identical binding sites on the IgG-Fc, single-domain SpA-B could, and indeed did, bind to IgG molecules with a stoichiometry of 2:1 (Fig 2A). SpA, however, although containing five IgG-binding domains, was found to bind principally to a single IgG molecule with a stoichiometry of 1:1 (Fig 2B). This suggests that full-length SpA may bind both IgG-Fc binding sites simultaneously.

**Figure 2.**
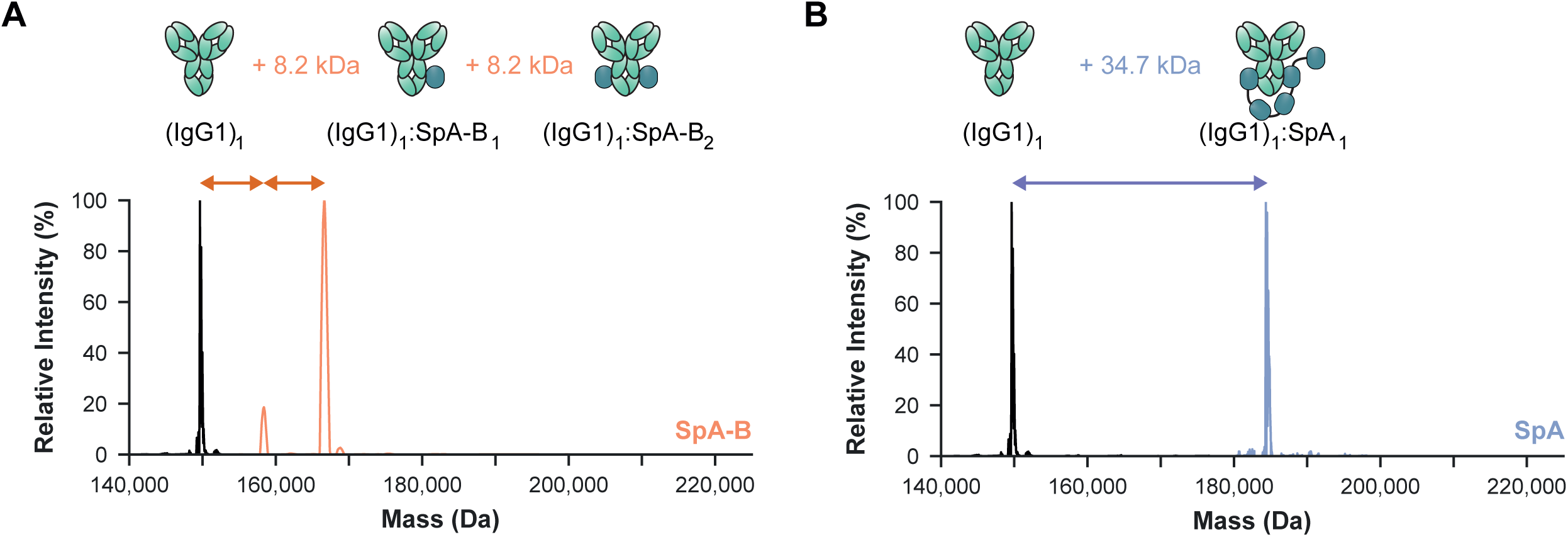
Single-domain SpA-B binds IgG1 with 2:1 stoichiometry, whereas five-domain SpA binds 1:1. A, B Deconvoluted native mass spectra show that the mass of IgG1 (black) is shifted when incubated with SpA-B (orange) (A) or SpA (blue) (B). This shift corresponds to binding of one or two copies of SpA-B to an IgG1 molecule and of only one SpA molecule to a single IgG1 molecule.

### SpA binding prevents IgG molecules from oligomerizing

Next, we evaluated whether SpA binding could reduce IgG hexamerization in solution. Although the formation of IgG oligomers is a surface phenomenon that requires IgG binding to antigenic surfaces, this process can be mimicked in solution by introducing three mutations in the Fc region of IgG that enhance Fc-Fc interactions (Diebolder *et al*, 2014; Wang *et al*, 2016). Combined mutation of residues E345**R** (Glu^345^→ Arg), E430**G** (Glu^430^→Gly) and S440**Y** (Ser^440^→Tyr) results in IgG-RGY, which readily forms hexamers in solution in a dynamic equilibrium (Diebolder *et al*, 2014; Wang *et al*, 2016). Here, we used native MS to investigate how SpA affects the monomer-hexamer equilibrium of IgG-RGY. As previously reported (Diebolder *et al*, 2014; Wang *et al*, 2016), native mass spectra of IgG1-RGY showed the presence of monomeric (denoted (IgG1)_1_) and hexameric ((IgG1)_6_) species, with intermediate states observed at lower abundance (Fig 3A and Table EV2). For IgG subclasses that bind SpA, the relative abundance of IgG oligomers was drastically reduced upon incubation with either SpA-B or SpA (Fig 3A, B and Table EV2). Under these conditions, we observed strong binding of SpA-B/SpA to IgG monomers, but not to the hexameric IgG species (Table EV2).

**Figure 3.**
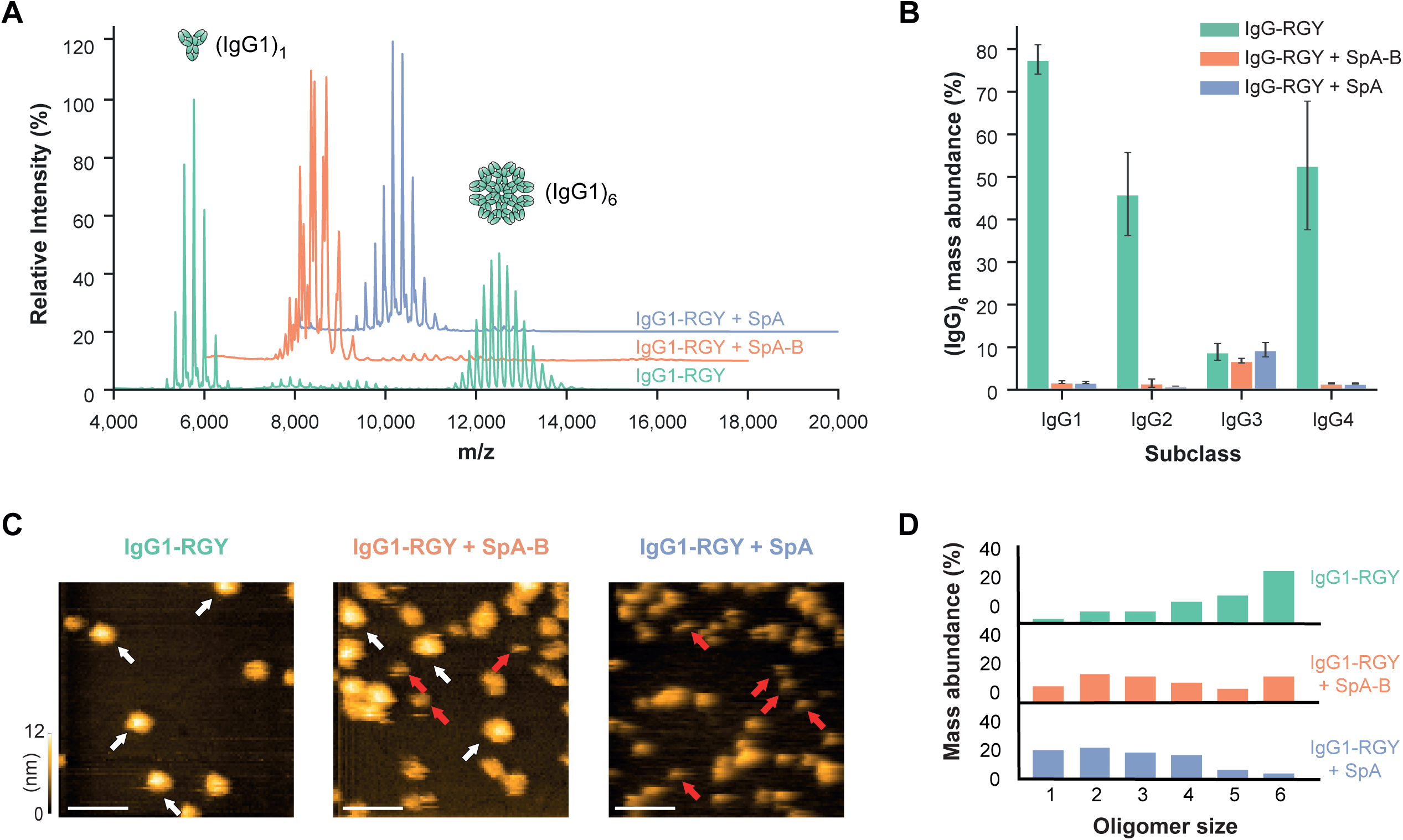
Binding of SpA-B or SpA blocks IgG-RGY monomers from assembling into higher-order oligomers. A Native mass spectra of IgG1-RGY in the absence (green) and presence of SpA-B (orange) or SpA (blue). B Hexamer relative mass abundance of IgG-RGY subclasses in the absence and presence of SpA-B or SpA, assessed by native MS. Error bars indicate the standard deviation over three replicate samples. C HS-AFM images of IgG1-RGY on DNP-lipid bilayers in the absence and presence of SpA-B or SpA, after pre-incubation in solution. White arrows: (IgG1)_6_. Red arrows: (IgG1)_1_. Scale bars: 100 nm. D IgG oligomer distribution of IgG1-RGY alone and in presence of SpA-B or SpA on DNP-lipid bilayers, after pre-incubation in solution. The histogram displays the fraction of IgGs constituting the respective oligomer species. Oligomer distribution was quantified by force-induced dissociation (IgG1-RGY: n=272; IgG1-RGY + SpA-B: n=697; IgG1-RGY + SpA: n=386).

Native MS measurements were corroborated by HS-AFM experiments on DNP-labeled-lipid-containing supported lipid bilayers (DNP-SLBs). Pre-incubation of the DNP-SLBs with anti-DNP antibody IgG1-RGY alone or in combination with SpA-B or SpA resulted in distinct distributions of differently sized IgG1-RGY oligomers (Fig 3C). Examination of oligomer size and overall IgG1 surface density by force-induced oligomer dissociation experiments (Strasser *et al*, 2019) allowed us to compile quantitative oligomer distributions (Fig 3D). Pre-incubation of anti-DNP IgG1-RGY with multi-domain SpA lead to a drastic reduction of higher-order IgG1-RGY oligomers on the DNP-SLB surface (Fig 3D) in agreement with our finding in the native MS experiments (Fig 3B). However, while the effect of both SpA constructs on the IgG1-RGY hexamer population in solution was comparable (Fig 3B), we observed only a ∼50% reduction of IgG1-RGY hexamers when IgG1-RGY was pre-incubated with the SpA-B domain on DNP-SLBs (Fig 3D). Altogether, these data suggest that SpA binds competitively to the Fc-Fc interaction site on IgG monomers, which effectively prevents IgGs from forming higher-order IgG oligomers.

### SpA prevents the formation of (IgG)_6_:C1q complexes in solution

Having demonstrated that SpA blocks IgG hexamerization in solution, we next verified whether SpA affects the formation of (IgG)_6_:C1q complexes. We used native MS to study the formation and behavior of these complexes in the presence and absence of SpA. In agreement with earlier data (Wang *et al*, 2016), native mass spectra of IgG1-RGY incubated with C1q showed clearly distinguishable species with masses corresponding to (IgG1)_1_, (IgG1)_6_ and (IgG1)_6_:C1q (Fig 4A and Table EV3). The experiments to study the effect of SpA were performed in two ways. When SpA or SpA-B was added to IgG1-RGY pre-incubated with C1q, the abundance of (IgG1)_6_:C1q complexes was strongly reduced (Fig 4A and Table EV3). Likewise, after pre-incubation of IgG1-RGY with the SpA constructs, the addition of C1q did not lead to detectible levels of (IgG1)_6_:C1q complexes. Thus, regardless of the order of mixing IgG1-RGY, C1q and SpA, we principally detected complexes of monomeric IgG bound to SpA constructs and free unbound C1q. Similar to IgG1-RGY, both SpA constructs also prevented the assembly of (IgG)_6_:C1q complexes for IgG2-RGY and IgG4-RGY (Fig 4B). Overall, these results suggest that SpA prevents the binding of C1q to IgG by blocking the formation of the hexameric IgG platform in solution.

**Figure 4.**
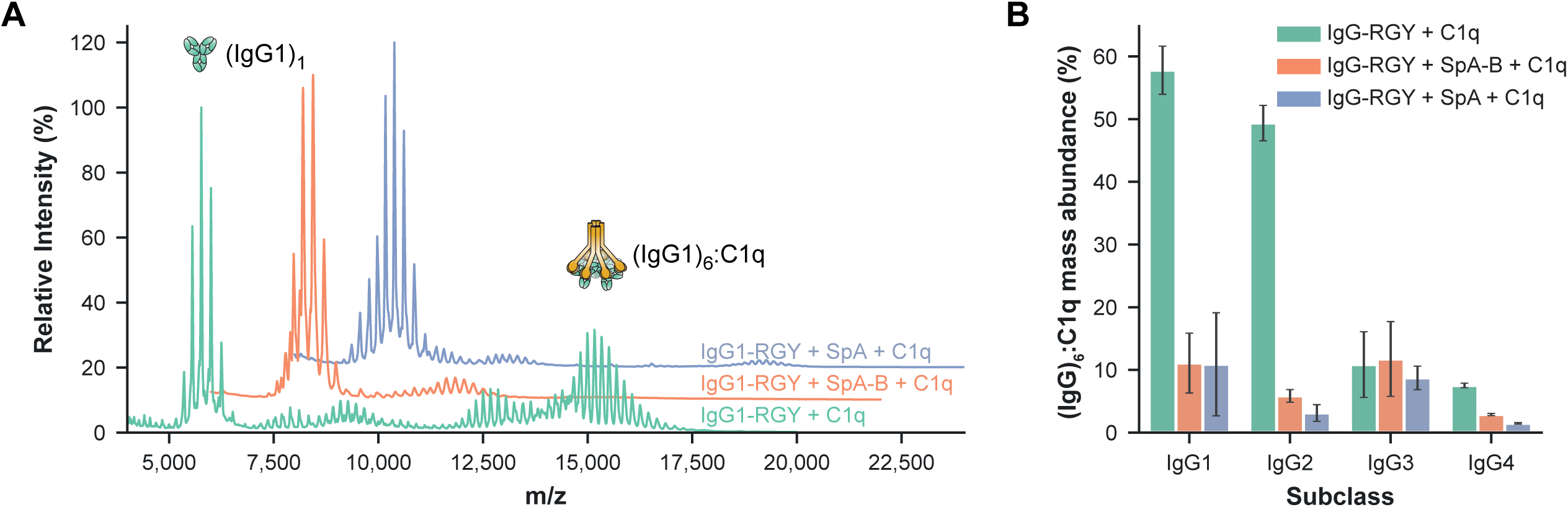
SpA-B and SpA prevent the assembly (IgG)_6_:C1q complexes in solution. A Native mass spectra of IgG1-RGY:C1q in the absence (green) and presence of SpA-B (orange) or SpA (blue). B The relative mass abundance of (IgG)_6_:C1q complexes in the absence and presence SpA-B or SpA, assessed by native MS. Error bars indicate the standard deviation over three replicate samples.

### SpA inhibits binding of C1q and C1 to antigen-bound IgGs on target surfaces

Next, we assessed how these observations for stabilized hexamers of mutant IgG-RGY in solution compare to the behavior of wild-type IgGs bound to antigenic surfaces. Since SpA and SpA-B block the formation of IgG hexamers through the same mechanism, we focused the following experiments on SpA-B. Using a quartz crystal microbalance (QCM) that we have previously used to quantitatively study binding and oligomerization of IgG1 to antigenic supported lipid bilayers (Strasser *et al*, 2020), we assessed the association of wild-type IgG1 to similar DNP-SLBs as used for HS-AFM experiments and followed the subsequent binding of C1q in the presence or absence of SpA-B (Fig 5A, B). Monitoring the resonance frequency change of the DNP-SLB-covered SiO_2_-coated quartz crystal, which is proportional to the change in bound mass, yields characteristic binding curves of anti-DNP IgG1-WT interacting with DNP-SLBs (Fig 5A-C, green time interval). As evident from the constancy of the sensorgrams in the following time interval (Fig 5A-C; grey time interval), the removal of IgG1-WT from the running buffer did not induce the dissociation of considerable amounts of IgG1-WT, thus reflecting stable binding. After having established equal amounts of surface-bound IgGs in all three experiments, we added C1q in the absence (Fig 5A, green time interval) and in the presence of SpA-B (Fig 5B, orange time interval) which resulted in distinct binding curves: while C1q robustly associated to IgG1-WT on our antigenic membranes (Fig 5A), simultaneous incubation of C1q with SpA-B strongly reduced the binding signal (Fig 5B). Addition of SpA-B alone (without C1q (Fig 5C)) resulted in strong association of SpA-B to IgG1-WT with no detectable dissociation when SpA-B was removed from the running buffer indicative for a high-affinity interaction. Notably, the remaining frequency shift (and thus the associated mass) at the end of the dissociation phase equaled the corresponding frequency shift for the C1q + SpA-B mixture, suggesting that the presence of SpA-B effectively precludes stable C1q binding (Fig 5B).

**Figure 5.**
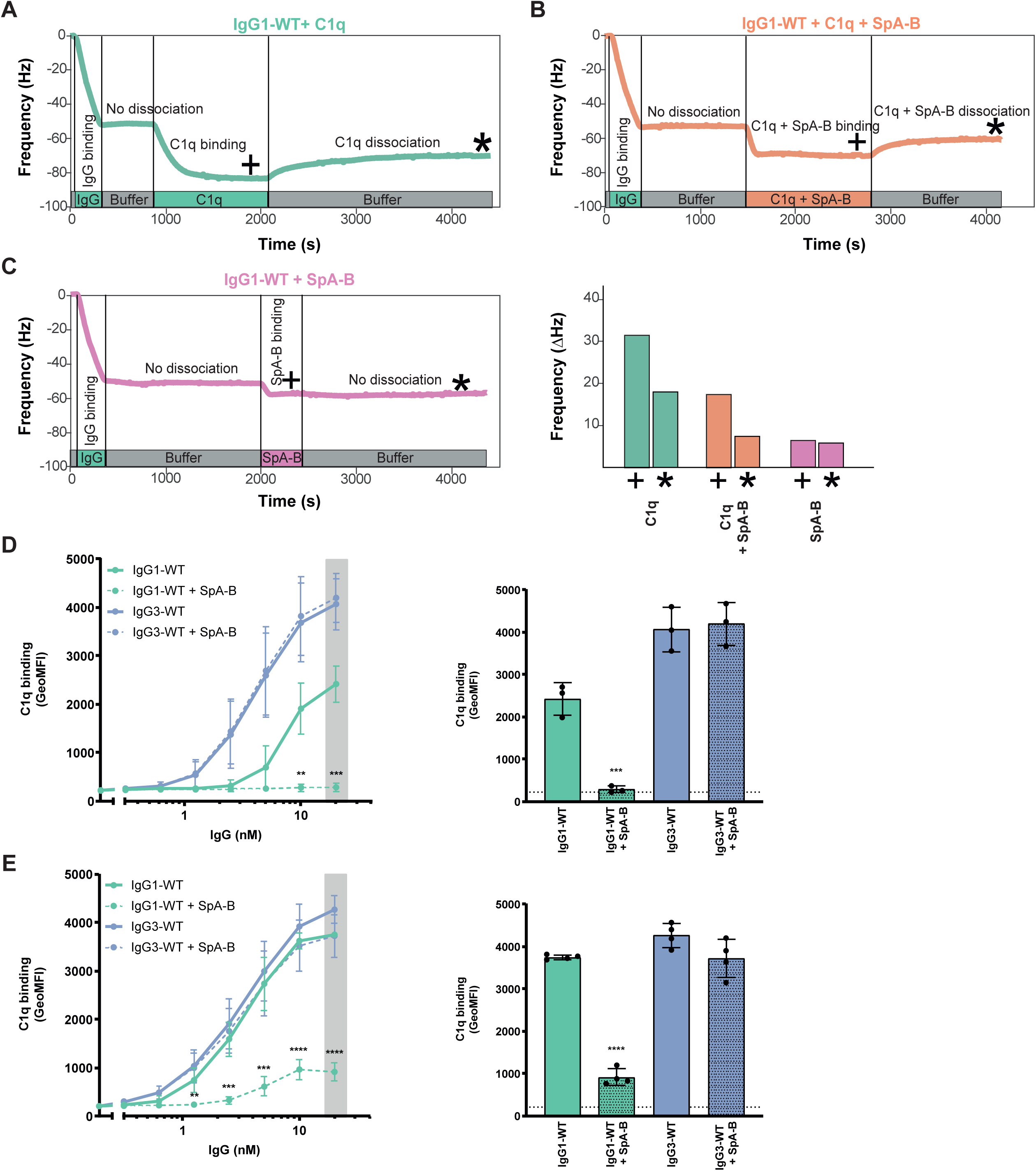
SpA-B inhibits binding of C1q to antigen-bound IgGs on target surfaces. A, B, C QCM sensorgrams of C1q alone (A), C1q and SpA-B (B), or SpA-B alone (C) binding to anti-DNP wild-type IgG1 antibodies (IgG1-WT) bound to DNP-decorated supported lipid bilayers. Binding of C1q (14 nM) was followed in real-time in the absence (A) or presence of SpA-B (B). Binding of SpA-B alone was monitored in a similar experiment without C1q (C). Bars represent the respective equilibrium level (+) and the level at the end of the dissociation phase (*). D, E C1q binding on IgG1-WT and IgG3-WT bound to DNP-coated beads, after incubation of C1q (D) or C1 complex (E) in the absence (solid lines) or presence (dotted lines) of SpA-B, detected with FITC-conjugated rabbit F(ab’)_2_ anti-human C1q by flow-cytometry. Bars represent the same data for the 20 nM IgG concentration only and the black dotted line shows the background fluorescence from beads that were not incubated with IgG. Data are presented as geometric means ± SD of three (D) or four (E) independent experiments. Statistical analysis was performed using an unpaired two-tailed t-test to compare buffer and SpA-B conditions and showed when significant as **P ≤ 0.01, ***P≤ 0.001 and ****P ≤ 0.0001.

QCM observations were corroborated by flow cytometry analyses, in which DNP-coated beads (Zwarthoff *et al*, 2018) were first labeled with anti-DNP IgG1-WT, or IgG3-WT as control. As expected, the presence of SpA-B inhibited C1q binding to IgG1-WT labeled beads, but binding to IgG3-WT was not altered (Fig 5D). As a control, we confirm that binding of IgG1-WT and IgG3-WT to DNP-coated beads was similar (Fig EV4A) while SpA-B could only bind to IgG1-WT-loaded target antigen (Fig EV4B). Interestingly, SpA-B also blocked C1q binding when incubated after IgG1-WT:C1q complexes were formed (Fig EV4C). Because the classical complement pathway is initiated by C1q that is in complex with C1r and C1s proteases, we repeated the same experiments using C1qr_2_s_2_ complexes instead of C1q alone. Also for this fully assembled C1 complex, we observed that SpA-B strongly reduced C1q binding to DNP-bound IgG1-WT and that its presence did not affect C1q binding to IgG3-WT (Fig 5E and EV4D). Finally, we confirmed that similar to SpA-B, SpA also largely reduces C1q binding to surface-bound IgGs, both on bead surfaces (Fig EV5A) and lipid membranes (Fig EV5B). Altogether, these data show that both SpA constructs efficiently block C1q binding to surface-bound antibodies of the IgG1 subclass.

### SpA inhibits C1q binding and complement activation on *S. aureus*

Finally, we determined whether SpA could affect antibody-dependent complement activation on the surface of *S. aureus*. To prevent cell-surface SpA (and Sbi, another Ig-binding protein of *S. aureus* (Zhang *et al*, 1998)) from binding the antibody’s Fc region, we here used a strain devoid of SpA and Sbi (NewmanΔ*spa/sbi*). NewmanΔ*spa/sbi* was labeled with a human monoclonal antibody directed against wall teichoic acid (WTA) (Lehar *et al*, 2015), a highly abundant anionic glycopolymer that is covalently anchored to the peptidoglycan layer (Brown *et al*, 2013). Although the anti-WTA antibody (clone 4497) belongs to VH3-type family (Fong *et al*, 2018), it lacks binding properties via its Fab region to SpA (Fig EV2C).

To measure C1q binding and downstream complement activation on the bacterial surface, we incubated IgG-labeled bacteria with human serum in combination with buffer, SpA-B or SpA. Because serum not only contains complement proteins but also many different antibodies (including IgGs that do not bind SpA (IgG3) and antibodies that bind SpA via their Fab region (VH3-type family Igs)), we used serum depleted from naturally occurring antibodies (ΔIgG/IgM serum). This way, we could exclusively determine the effect of SpA on complement activation by monoclonal anti-WTA antibodies. In agreement with the results from experiments using DNP-beads, we observed that both SpA constructs strongly reduced C1q binding on *S. aureus* labeled with anti-WTA IgG1 antibodies (Fig 6A). When the bacteria were labeled with anti-WTA IgG3, C1q binding was not inhibited by the presence of SpA-B or SpA (Fig 6B).

**Figure 6.**
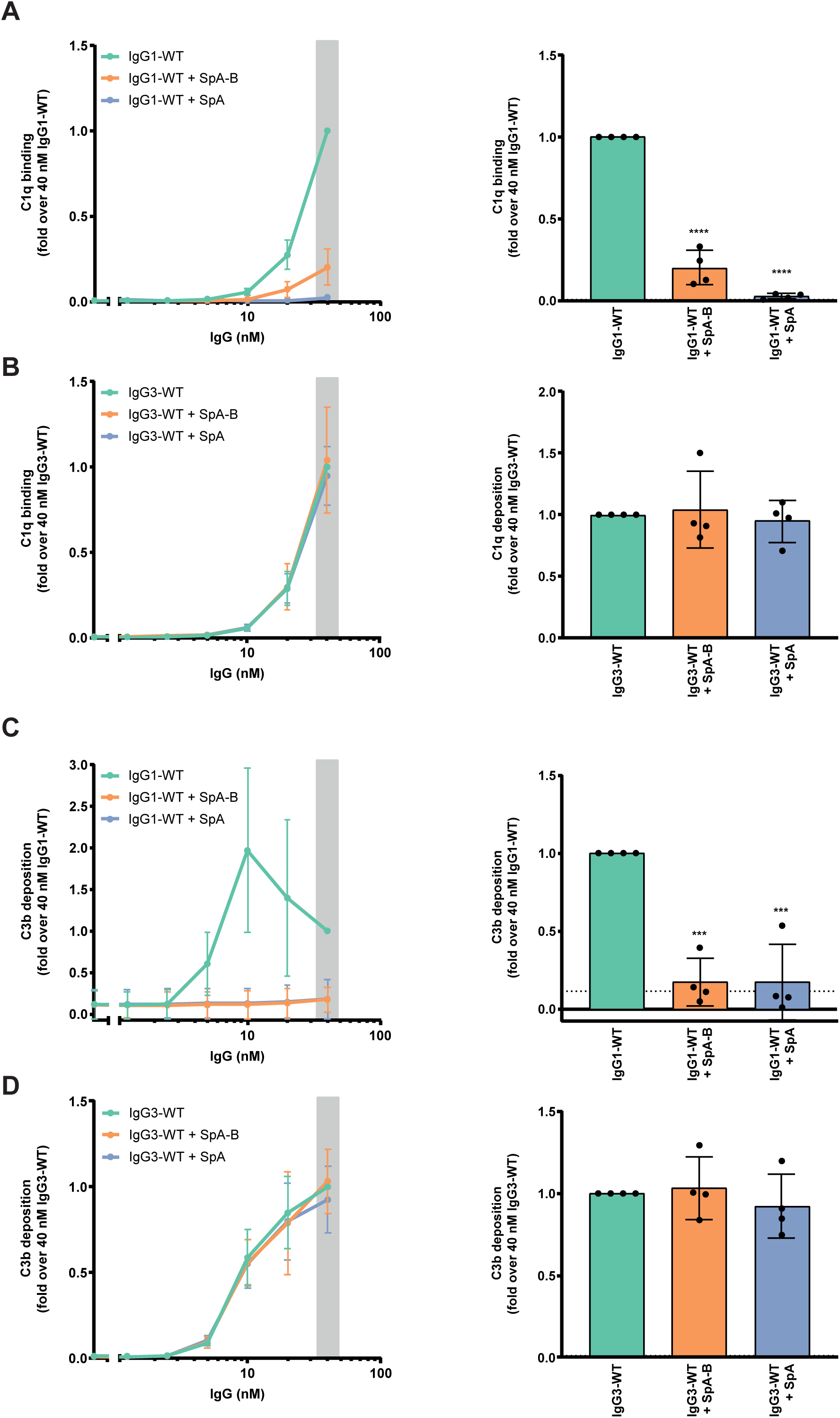
SpA decreases IgG-mediated C1q binding and downstream complement on *S. aureus*. A, B C1q binding on anti-WTA wild-type IgG1 (IgG1-WT) (A) or IgG3 antibodies (IgG3-WT) (B) bound to NewmanΔ*spa*/*sbi* surface, after incubation of bacteria with 1% ΔIgG/IgM human serum, in the absence (green) or presence of SpA-B (orange) or SpA (blue), detected with FITC-conjugated rabbit F(ab’)_2_ anti-human C1q by flow cytometry. C, D C3b deposition on NewmanΔ*spa*/*sbi* surface after incubation of bacteria with IgG1-WT (A) or IgG3-WT (B), 1% ΔIgG/IgM human serum and buffer (green), SpA-B (orange) or SpA (blue), detected with a monoclonal murine anti-human C3d antibody by flow cytometry. Data information: In (A-D), data are presented as fold change over 40 nM concentration of IgG control ± SD of at least three independent experiments. Bars represent the same data for the 40 nM IgG concentration only and the black dotted line shows the background fluorescence from bacteria that were not incubated with IgG. Statistical analysis was performed using a one-way ANOVA to compare buffer condition with SpA-B and SpA conditions and displayed when significant as ***P ≤ 0.001 or ****P ≤ 0.0001.

To determine whether the inhibition of C1q binding leads to a downstream inhibition of the complement cascade, we also measured the deposition of C4b and C3b. The binding of C1q to target-bound antibodies activates its attached C1r/C1s proteases that cleave C4 and C2, and generate the C3 convertase (C4b2a) (Law & Dodds, 2008). Subsequently, the covalently attached C3 convertase catalyzes the deposition of C3b on the target surface (Law & Dodds, 2008). Upon incubation of IgG1-labeled *S. aureus* with human (Ig-depleted) serum and SpA-B or SpA, surface deposition of C4b and C3b was completely abolished (Fig EV6A and 6C). As anticipated, C4b and C3b deposition on NewmanΔ*spa/sbi* labeled with anti-WTA IgG3 antibodies remained unchanged in the presence of SpA-B or SpA (Fig EV6B and 6D). Altogether, these findings demonstrate that SpA effectively prevents C1q recruitment and downstream activation of the complement cascade on *S. aureus*.

## Discussion

Antibody-dependent complement activation is an important immunological mechanism to accelerate bacterial killing (Lu *et al*, 2018). To effectively trigger complement, antibodies should bind bacterial cells and subsequently form oligomeric IgG clusters that recruit C1 (Diebolder *et al*, 2014; Strasser *et al*, 2019). In this paper, we identify SpA from *S. aureus* as the first example of a bacterial immune evasion molecule that specifically blocks antibody clustering by inhibiting IgG Fc-Fc contacts. These findings are of relevance to the basic pathophysiology of staphylococcal infections, but also to the development of immune therapies against *S. aureus*. Furthermore, SpA could be used as a tool to better understand the role of IgG clustering in various disease processes where antibodies and complement are involved.

Our study demonstrates that, by binding to the Fc region of IgG, soluble SpA blocks IgG hexamerization, which results in inhibition of C1q recruitment and downstream complement activation on the *S. aureus* surface. Although the interference of SpA with the classical complement pathway has been noted before, the exact molecular mechanism remained elusive and there appeared to be a discrepancy about its role as an activator (Sjöquist & Stålenheim, 1969, 1970; Kronvall & Gewurz, 1970; Stålenheim & Malmheden-Eriksson, 1971; Stålenheim & Castensson, 1971; Stalenheim, 1971; Stålenheim *et al*, 1973) or inhibitor (Kronvall & Gewurz, 1970; Stålenheim *et al*, 1973) of the complement system. Thanks to recent insights into IgG oligomerization (Diebolder *et al*, 2014) and more advanced methods to directly visualize this process, we have been able to unravel the mechanism of SpA-dependent complement inhibition in highly purified conditions. The binding of SpA to the Fc region of monomeric, but not hexameric, IgG-RGY species suggests that SpA exclusively binds to free, non-occupied Fc regions. Thus, by binding to monomeric IgGs, SpA likely prevents the transition to the hexameric state.

Since *S. aureus* strains produce SpA with five or four Ig-binding domains (Baum *et al*, 2009), we here compared the complement inhibitory activity of SpA with five Ig-binding domains versus a single SpA-B domain. AFM experiments suggest that SpA more effectively blocks hexamerization of IgG molecules on antigenic membranes than SpA-B. Because multi-domain SpA is likely to bind IgG bivalently, it dissociates at a lower rate than monovalently bound SpA-B. The resulting binding affinity advantage of SpA over SpA-B would allow it to compete more effectively with the IgG Fc-Fc interactions. Indeed, multi-domain SpA was more effective in inhibiting C1q binding to target-bound IgGs than SpA-B, which is consistent with the notion that multivalent SpA molecules have a complement inhibitory advantage over a single SpA domain. However, upon measuring downstream complement activation, the inhibitory advantage of SpA over a single SpA domain was less evident. We speculate that the requirement for multiple domains may be even more important for complement inhibition by cell-anchored SpA. Since the majority of SpA produced by *S. aureus* is anchored to the bacterial cell wall, multiple SpA domains may be needed to provide the molecule with sufficient length and flexibility to bind Fc domains of bacterium-bound IgG. Although the majority of SpA is cell-anchored, 6.5% to 15% is secreted before sorting (O’Halloran *et al*, 2015) or released from the cell wall after enzymatic cleavage by LytM (Becker *et al*, 2014). We observed that the concentration of recombinant multi-domain SpA needed to decrease C1q deposition on an antigenic surface is lower than the amount reported to be secreted by Newman or USA300 strains *in vitro* (Hoppenbrouwers *et al*, 2018).

Besides SpA, there are other bacterial proteins that bind to Ig (Sidorin & Solov’Eva, 2011). The best known are Protein G (from Group C and G streptococci (Björck & Kronvall, 1984; Reis *et al*, 1984)), M and M-like proteins (Group A streptococci (Heath *et al*, 1990; Stenberg *et al*, 1992; Boyle *et al*, 1994)), Protein L (*Peptostreptococcus magnus* (Bjorck, 1988)), Sbi and SSL10 (*S. aureus* (Zhang *et al*, 1998; Itoh *et al*, 2010)). Protein G, Protein M, M-like proteins, and Sbi all bind the Fc domain of IgGs in the vicinity of the SpA binding site (Atkins *et al*, 2008; Sauer-eriksson *et al*, 1995; Frick *et al*, 1992), which suggests that these IgG-Fc binding proteins may be able to block IgG hexamerization as well.

Overall, our data provide insights that are crucial for the development of effective immune therapies against *S. aureus*. It is well-recognized that both neutrophils and complement play an essential role in the killing of *S. aureus*. While neutrophils can engulf *S. aureus* directly, earlier work from our group has demonstrated that the decoration of bacteria with C3-derived opsonins strongly enhances effective phagocytosis (Rooijakkers *et al*, 2005). Therefore, the identification of antibacterial antibodies with strong complement-activating potential provides an interesting approach to boost the host immune system and prevent or treat these infections. From this study, it is now clear that such approaches can only be successful if we take the SpA-dependent antibody modulation into account. Although most active and passive immunization approaches to develop or induce antibodies targeting *S. aureus* surface components (e.g. capsule polysaccharide, LTA and various surface adhesins) have failed at the clinical trial stage (Sause *et al*, 2016), we propose that such strategies could be more effective when also blocking the effect of SpA. In this context, our comparison of different IgG subclasses suggests that monoclonal antibodies targeting surface components of *S. aureus* should be developed as IgG3 (or variants thereof) that are not targeted by SpA (Inganäs, 1981). Alternatively, any amino acid modification in IgG1 or IgG2 subclasses that prevents SpA binding to IgG, but not IgG hexamerization, would be of value. We also propose that the use of SpA as a vaccine antigen or of monoclonal antibodies directed against Fc-binding domains of SpA might be needed to prevent the anti-complement effect of SpA and therefore, increase the chances of bacterial clearance.

Furthermore, SpA could also be used as a research tool to specifically examine the role of IgG hexamerization in various disease processes. For instance, SpA or a single domain of SpA (mutated to only bind IgG-Fc region) could be used to study whether antibodies induced during an infection require Fc-Fc contacts to induce complement activation on the invading pathogen. In fact, our data with IgG1 antibodies against WTA suggest that naturally induced antibodies against *S. aureus* indeed require Fc-Fc contacts to induce complement activation on the surface of bacteria. Although we used monoclonal antibodies, it is well-recognized that IgG1 antibodies against WTA are produced during an *S. aureus* infection *in vivo* (Jung *et al*, 2012; Lee *et al*, 2015; Kurokawa *et al*, 2016; van Dalen *et al*, 2019). Also, given that excessive activation of complement has been associated with clinical manifestation of several autoimmune diseases, SpA could be used to study whether autoreactive antibodies induce IgG clustering on altered host cells. Finally, SpA, or molecules targeting the SpA-binding site in IgG, has the potential to block unwanted antibody responses. Indeed, the therapeutic potential of SpA has been tested for treatment of autoimmune disorders (Eftimiadi *et al*, 2017; Kapur *et al*, 2018; Bernton *et al*, 2014; Viau & Zouali, 2005), although the reasoning for the use of SpA was based on its effectiveness as a B cell superantigen.

In conclusion, the identification of SpA as the first biological inhibitor of IgG hexamerization will increase our understanding of antibody-dependent immunological mechanisms and may help to accelerate the development of immune interventions in infection and inflammation.

## Materials and methods

### Production and purification of human monoclonal antibodies

Monoclonal antibodies against DNP (DNP-G2a2) used for MS, HS-AFM and QCM experiments were recombinantly expressed as wild-type and hexamer-forming RGY mutant (Diebolder *et al*, 2014) IgG1, IgG2, IgG3 and IgG4 and obtained from Genmab (Utrecht, the Netherlands) (White *et al*, 1996; Ugurlar *et al*, 2018).

For the other experiments of this study, we used human monoclonal antibodies produced recombinantly in human Expi293F cells (Life Technologies) as described before (Fang *et al*, 2017), with minor modifications. Briefly, gBlocks (Integrated DNA technologies, IDT), containing codon-optimized variable heavy and light chain (VH and VL) sequences with an upstream KOZAK and HAVT20 signal peptide, were cloned into homemade pcDNA34 vectors, upstream the IgG heavy and kappa light chain constant regions, respectively, using Gibson assembly (New England Biolabs). The VH and VL sequences of the antibodies were derived from previously reported antibodies anti-DNP (DNP-G2a2) (Gonzalez *et al*, 2003) and anti-WTA GlcNAc-β-4497 (patent WO/2014/193722) (Lehar *et al*, 2015) (Table EV4). Transfection of EXPI293F cells was performed using PEI (Polyethylenimine HCl MAX; Polysciences). After 4 to 6 days of transfection, IgG1, IgG2 and IgG4 antibodies were isolated from cell supernatants using a HiTrap Protein A High Performance column (GE Healthcare), whereas IgG3 antibodies were isolated with a HiTap Protein G High Performance column (GE Healthcare). Antibodies were dialyzed in PBS, overnight at 4 °C, and filter-sterilized through 0.22 µm Spin-X filters. Antibodies were analyzed by size exclusion chromatography (GE Healthcare) and monomeric fractions were isolated in case of aggregation levels >5%. The concentration of the antibodies was determined by measurement of the absorbance at 280 nm and antibodies were stored at −20 °C until use. The anti-Hla (MEDI4893; patent WO/2017/075188A2), which served as a control, was a gift from Dr Alexey Ruzin, MedImmune (AstraZeneca).

### Antibody cleavage

Anti-DNP IgG1 Fc and F(ab’)_2_ molecules were generated by overnight digestion at 37 °C using 0.25 U/µg FabRICATOR IdeS (Genovis AB).

FITC-conjugated rabbit F(ab’)_2_ anti-human C1q was obtained by digestion of FITC-conjugated rabbit anti-human C1q (Dako) using 1 U/ µg of recombinant His-tagged IdeS protease. After incubation of antibody with IdeS for 2 h at 37 °C in the dark, F(ab’)_2_ fragment was purified though HiTrap Protein A High Performance column (GE Healthcare) and HisTrap High Performance column (GE Healthcare).

### Cloning, expression and purification of SpA-B constructs

Codon-optimized gBlocks (IDT) for wild-type B domain of SpA (SpA-B), SpA-B lacking Fc-binding properties (SpA-B^KK^; Q9K and Q10K mutations) and SpA-B lacking Fab-binding properties (SpA-B^AA^; D36A and D37A mutations) (Table EV5), containing a C-terminal LPETG sortagging sequence, were cloned into a modified pRSET-C-HIS vector by Gibson assembly. Recombinant proteins, containing a C-terminal LPETGG-AAA-HHHHHH tag, were generated in E. coli BL21(DE3) and were isolated under native purification conditions using a 5 HisTrap High Performance Column (GE Healthcare) with imidazole gradient (10-250 mM; Sigma-Aldrich). SpA-B constructs were dialyzed in 50 mM Tris 300 mM NaCl pH 8.0, overnight at 4 °C, and stored at −20 °C until use.

### Native mass spectrometry

Native MS analyses were performed on a modified LCT time-of-flight instrument (Waters) or a standard Exactive Plus EMR Orbitrap instrument (ThermoFisher Scientific). Before analysis, buffers were exchanged to 150 mM ammonium acetate (pH 7.5) through six consecutive dilution and concentration steps at 4 °C using Amicon Ultra centrifugal filters with a 3 kDa or 10 kDa molecular weight cutoff (Merck). Protein complexes were assembled by mixing the subcomponents at the desired molar ratios, followed by incubation at RT for at least 30 min. For experiments studying the effect of SpA constructs, the incubation step with SpA proceeded for at least 3 h at 37 °C due to the relatively slow disassembly rate of the IgG-RGY hexamers. IgG-RGY hexamers were measured at a total IgG concentration of 2 µM in the presence or absence of 10 µM SpA-B or SpA (ProSpec). For measurements of (IgG)_6_:C1q complexes, 0.5 µM C1q (Complement Technology) was used. Samples were loaded into gold-coated borosilicate capillaries (prepared in-house) for direct infusion from a static nano-ESI source. Relative abundances of protein complexes were determined using an in-house script that sums and compares ion intensities of the different species, similar to a previously described method (Wang *et al*, 2012). Deconvoluted mass spectra were generated by Bayesian deconvolution using UniDec (Marty *et al*, 2015).

### Antibody labeling

IgG labeling was performed by incubation of antibodies with Alexa Fluor^647^ NHS Ester (Succinimidyl Ester; ThermoFisher Scientific). In detail, 1 µL of 10 mg/mL Alexa Fluor^647^ NHS Ester was added to 100 µl of 1 mg/mL antibody in PBS with 0.1 M sodium bicarbonate buffer. After 2 h incubation at RT in the dark, labeled antibodies were separated from the free probe by use of Pierce desalting spin column 0.5 mL (ThermoFisher Scientific), according to the manufacturer’s manual. Subsequently, absorbance was measured at 280 nm for protein and 647 nm for the probe, using a Nanodrop.

### Binding of antibodies to beads coated with SpA constructs

Dynabeads His-Tag Isolation & Pulldown (Invitrogen) were washed in PBS-TH (PBS, 0.05% (v/v) Tween-20 and 0.5% human serum albumin (HSA)). 50 µL of 30 µg/mL his-tagged SpA-B, SpA-B^KK^ or SpA-B^AA^ was mixed with 1 µL beads for 30 min at 4 °C, shaking (± 700 rpm). After two washes with PBS-TH, 0.05 µL beads were incubated with 2 µg/mL anti-DNP IgG1, anti-DNP IgG3 or anti-WTA IgG1 in 30 µL, for 30 min at 4 °C, shaking (± 700 rpm). A VH3-type antibody, anti-Hla IgG1, was included as a control for Fab binding to both SpA-B and SpA-B^KK^. Anti-DNP IgG3 was included as a negative control. All antibodies were also incubated with Dynabeads Protein G (Invitrogen) that served as reference binding for all antibodies, including IgG3. Antibody binding was detected using directly labeled IgG or Alexa Fluor^647^-conjugated goat F(ab’)_2_ anti-human kappa (Southern Biotech). Samples were measured using flow cytometry (BD FACSVerse) and data, based on a single bead population, were analyzed using FlowJo software. Mean fluorescence values were expressed relative to the control binding to Protein-G beads for each antibody.

### Enzyme-Linked Immunosorbent Assay (ELISA)

MaxiSorp plates (Nunc) were coated with 3 ug/mL SpA-B or SpA in 0.1 M sodium carbonate, overnight at 4 °C. The day after, plates were washed three times with PBS-T (PBS, 0.05% (v/v) Tween-20) and blocked with 4% bovine serum albumin (BSA) in PBS-T, for 1 h at 37 °C. The following incubations were performed for 1 h at 37 °C followed by 3 washes with PBS-T. Five-fold serial dilutions (starting from 20 nM) of human monoclonal anti-DNP IgG1, IgG2, IgG3 and IgG4 in 1% BSA in PBS-T were added to the wells. Bound antibodies were detected with horseradish peroxidase (HRP)-conjugated goat F(ab’)_2_ anti-human kappa (Southern Biotech) in 1% BSA in PBS-T and Tetramethylbenzidine (TMB) as substrate. The reaction was stopped with 1N sulfuric acid and absorbance was measured at 450 nm in the iMark™ Microplate Absorbance Reader (Bio-Rad).

### DNP labeled liposomes

DNP-labeled liposomes consisting of 1,2-dipalmitoyl-sn-glycero-3-phosphocholine (DPPC), 1,2-dipalmitoyl-sn-glycero-3-phosphoethanolamine (DPPE) and 1,2-dipalmitoyl-sn-glycero-3-phosphoethanolamine-N-[6-[(2,4-dinitrophenyl)amino]hexanoyl] (DNP-cap-DPPE) were used to generate supported lipid bilayers (SLBs) on mica and SiO_2_ substrates. The lipids were purchased from Avanti Polar Lipids, mixed at a ratio of DPPC:DPPE:DNP-cap-DPPE = 90:5:5 (molar ratio), and dissolved in a 2:1 mixture of chloroform and methanol. After the solvents were rotary-evaporated for 30 min, the lipids were again dissolved in chloroform, which was then rotary-evaporated for 30 min. Drying was completed at a high vacuum pump for 2 h. The lipids were dissolved in 500 µL Milli-Q H_2_O while immersed in a water bath at 60 °C, flooded with argon, and sonicated for 3 min at 60 °C to create small unilamellar vesicles. These were diluted to 2 mg/mL in buffer #1 (10 mM HEPES, 150 mM NaCl, 2 mM CaCl_2_, pH 7.4) and frozen for storage using liquid N_2_.

### High-speed atomic force microscopy

HS-AFM (RIBM, Japan) was conducted in tapping mode at RT in buffer, with free amplitudes of 1.5 - 2.5 nm and amplitude set points larger than 90%. Silicon nitride cantilevers with electron-beam deposited tips (USC-F1.2-k0.15, Nanoworld AG), nominal spring constants of 0.15 N m^-1^, resonance frequencies around 500 kHz, and a quality factor of approx. 2 in liquids were used. Imaging was performed in buffer #1. All IgGs were diluted and incubated in the same buffer.

DNP labeled supported lipid bilayers for HS-AFM were prepared on muscovite mica. The liposomes were incubated on the freshly cleaved surface (500 µg mL^-1^ in buffer #1), placed in a humidity chamber to prevent evaporation, and heated to 60 °C for 30 min. Then the temperature was gradually cooled down to RT within 30 min, followed by exchanging the solution with buffer #1. After 10 min of equilibration at RT and 15 more buffer exchanges, the SLB was ready for imaging. In order to passivate any exposed mica, SLBs were incubated with 333 nM IgG1-b12 (irrelevant human IgG1 control antibody against HIV-1 gp120) (Burton *et al*, 1994) for 10 min before the molecules of interest were added.

### IgG oligomer statistics on DNP-SLBs

For experiments studying the effect of SpA constructs, the incubation step with SpA proceeded for at least 2 h at 37 °C. IgG-RGY was used at a total IgG concentration of 2 µM in the presence or absence of 20 µM SpA-B or SpA. IgG oligomer distributions were then obtained by incubating DNP-SLBs with 33.3 nM of the respective IgG variant for 5 min, and analyzed in a two-step process: Individual particle dimensions were determined by HS-AFM, and the respective oligomeric states were further confirmed by observing their decay pattern determined in subsequent forced dissociation experiments as described (Strasser *et al*, 2019). In brief, molecules are scanned in a nondisrupting manner to gauge their number, height, and shape. Subsequently, the scanning force exerted by the HS-AFM cantilever tip is increased to dissociate oligomers into their constituent IgGs. Geometric parameters and oligomer decay patterns are combined to assign each IgG assembly its oligomeric state.

### Quartz crystal microbalance (QCM)

QCM experiments were done using a two-channel QCM-I system from MicroVacuum Ltd. (Budapest, Hungary). AT-cut SiO_2_-coated quartz crystals with a diameter of 14.0 mm and a resonance frequency of 5 MHz were used (Quartz Pro AB, Jarfalla, Sweden). All sensorgrams were recorded on the first, third and fifth harmonic frequencies. The data shown relates to the third harmonic. Before each set of experiments, the SiO_2_-coated crystals were cleaned by immersion in 2% SDS for 30 min, followed by thorough rinsing with Milli-Q H_2_O. The chips were dried in a gentle stream of N_2_ and oxidized using air plasma (4 min at 80 W), after which they were mounted in the measurement chamber. The sensor surface was further cleaned by a flow of 2% SDS at 250 µL min^-1^ for 5 min, followed by Milli-Q H_2_O at 250 µL min^-1^ for 5 min directly before the measurement. Finally, IgG-free buffer was injected for equilibration at 50 µL min^-1^. All subsequent injections were performed at 50 µL min^-1^. To generate DNP-SLBs on QCM chips, the DPPC:DPPE:DNP-cap-DPPE liposome stock solution was heated to 60 °C for 30 min, and slowly cooled down to RT within 30 min. The solution was ready for injection after dilution to 200 µg/mL with buffer #1. DNP-SLB formation was typically complete after 30 min at 50 µL min^-1^, after which the flow medium was changed to buffer #1 for equilibration.

### DNP-coated beads assays

Dynabeads M-270 Streptavidin beads (Invitrogen) were washed in PBS-TH and incubated, 100 times diluted, with 1 µg/mL of biotinylated 2,4-dinitrophenol (DNP-PEG2-GSGSGSGK(Biotin)-NH2, Pepscan Therapeutics B.V.) in PBS-TH for 30 min at 4 °C, shaking (± 700 rpm). For each condition, 0.5 µL of beads were used (∼3×10^5^ beads/condition). After two washes with PBS-TH, DNP-coated beads were incubated with 20 nM anti-DNP IgG or 2-fold serial dilutions of anti-DNP IgG (starting from 20 nM IgG), for 30 min at 4 °C, shaking (± 700 rpm). The following incubations steps were performed under shaking conditions (± 700 rpm) for 30 min at 4 °C unless stated otherwise. Additionally, after each incubation, beads were washed twice with PBS-TH or VBS-TH (Veronal Buffered Saline pH 7.4, 0.5 mM CaCl_2_, 0.25 mM MgCl_2_, 0.05% (v/v) Tween-20, 0.5% HSA), dependent on buffer used in the following incubation step.

For antibody binding detection, IgG-bound DNP-beads were next incubated with 1 µg/mL Alexa Fluor^647^-conjugated goat F(ab’)_2_ anti-human kappa (Southern Biotech) in PBS-TH.

For SpA binding detection, IgG-bound DNP-coated beads were first incubated with 200 nM of recombinant His-tagged SpA-B or SpA (ProSpec) in PBS-TH. Subsequently, beads were incubated with 1 µg/mL chicken anti-HexaHistidine antibody (Nordic) in PBS-TH, followed by incubation with R-Phycoerythrin (PE)-conjugated donkey F(ab’)_2_ anti-chicken (Jackson ImmunoResearch) diluted 1/500 in PBS-TH.

For C1q binding experiments, IgG-bound DNP-coated beads were incubated with 1.3 nM of purified C1 (Complement Technology) or C1q (Complement Technology) alone, in combination with, or followed by incubation of 200 nM or 4-fold dilutions (starting from 1 µM) of recombinant SpA-B or SpA in VBS-TH, for 30 min at 37 °C. C1q was detected by use of 4 µg/mL FITC-conjugated rabbit F(ab’)_2_ anti-human C1q. The binding of antibody, SpA and C1q to the beads were detected using flow cytometry (BD FACSVerse) and data were analyzed based on single bead population using FlowJo software.

### Depletion of IgG and IgM from human serum

Human serum was pooled from 20 healthy donors. Informed consent was obtained from all subjects in accordance with the Declaration of Helsinki. Approval from the Medical Ethics Committee of the University Medical Center Utrecht was obtained (METC protocol 07-125/C approved on March 1, 2010). Human serum was depleted of IgG and IgM as previously reported (Zwarthoff *et al*). Briefly, IgG and IgM were captured by affinity chromatography using HiTrap Protein G High Performance column (GE Healthcare) and CaptureSelect IgM Affinity Matrix (ThermoFisher Scientific), respectively. After depletion, antibodies and complement levels were quantified by ELISA and complement activity was measured by classical pathway and alternative pathway hemolytic assays. As IgG and IgM depletion results in partial co-depletion of C1q, ΔIgG/IgM serum was reconstituted with purified C1q (Complement Technology) to physiological levels (70 µg/mL).

### Complement deposition assays on *S. aureus* surface

*S. aureus* NewmanΔ*spa*/*sbi* strain was fluorescently labeled by transformation with the pCM29 plasmid, constitutively expressing mAmetrine under the regulation of the sarA promotor, as described before (Pang *et al*, 2010; Schenk & Laddaga, 1992; De Jong *et al*, 2017). Bacteria were grown overnight in Todd Hewitt broth (THB) plus 10 µg/mL chloramphenicol, diluted to an OD_600_ of 0.05 in fresh THB plus chloramphenicol and cultured until mid-log phase (OD_600_ = 0.5). Cells were collected, washed and resuspended in RPMI + 0.05% HSA, and aliquoted at −20 °C.

Similarly to Dynabeads assays, all incubation steps were performed under shaking conditions (± 700 rpm) for 30 min at 4 °C, followed by a single wash with RPMI-H by centrifugation. Bacteria (7.5×10^5^ CFU) were incubated with 2-fold titration (starting from 40 nM) IgG in RPMI-H followed by incubation with 1% ΔIgG/IgM serum in combination with buffer, 200 nM of SpA-B or SpA in RPMI-H, for 30 min at 37 °C. For C1q detection, bacteria were incubated with 0.5 µg/mL of chicken anti-human C1qA (Sigma-Aldrich), followed by incubation with PE-conjugated donkey F(ab’)_2_ anti-chicken (Jackson ImmunoResearch), diluted 1/500 in RPMI-H.

For C4b and C3b detection, 1 µg/mL of murine anti-C4d (Quidel) or anti-C3d (Quidel) antibody was incubated with bacteria, respectively. Subsequently, bacteria were incubated with FITC-conjugated goat F(ab’)_2_ anti-mouse (Dako), diluted 1/100 in RPMI-H.

After labelling, samples were fixed with 1% paraformaldehyde in RPMI-H and the binding of C1q, C4b and C3b to bacteria was detected using flow cytometry (BD FACSVerse). Data were analyzed using FlowJo software.

### Statistical analysis

Statistical analysis was performed with GraphPad Prism v.8.3 software, using unpaired two-tailed t-test or one-way ANOVA as indicated in the figure legends. At least three experimental replicates were performed to allow statistical analysis.

## Acknowledgements

This work was supported by the Netherlands Organization for Scientific Research (NWO) through a TTW-NACTAR Grant #16442 (to AJRH and SHMR), the European Union’s Horizon 2020 research programs H2020-MSCA-ITN (#675106, to JAGS and FB) and ERC Starting grant (#639209, to SHMR). MAB and AJRH further acknowledge funding for the large-scale proteomics facility, the Netherlands Proteomics Center, through the X-omics Road Map program (project 184.034.019) and the EU Horizon 2020 program INFRAIA project Epic-XS (Project 823839). AJRH and SHMR acknowledge the Utrecht University Molecular Immunology HUB. JP acknowledges support by the European Fund for Regional Development (EFRE, Regio 13) and the Federal State of Upper Austria. The authors thank Annette Stemerding for fruitful discussions.

## Author contributions

ARC, MAB, JS, FJB, GW, RNJ, JAGS, KPMK, JS, JP, AJRH and SHMR were involved in designing or conceiving the study. ARC, MAB, JS, CJCH, PCA and KPMK performed experiments and analyzed the data. RNJ generated atomic models of IgG hexamers and SpA. ARC, MAB, JS, KPMK, JP, AJRH and SHMR wrote the paper. FB, JAGS, JS, JP, AJRH and SHMR acquired the funding. KPMK, JP, AJRH and SHMR supervised the project. All authors critically revised the manuscript and approved it before submission.

## Conflict of interest

ARC participated in a postgraduate studentship program at GSK. FJB, JAGS, KPMK, JS and SHMR are co-inventor on a patent describing antibody therapies against *S. aureus*. FJB, RNJ and JS are Genmab employees. FB is an employee of GSK group of companies and is co-inventor on patents on *S. aureus* vaccine candidates.

## Expanded View

**Figure EV1.**
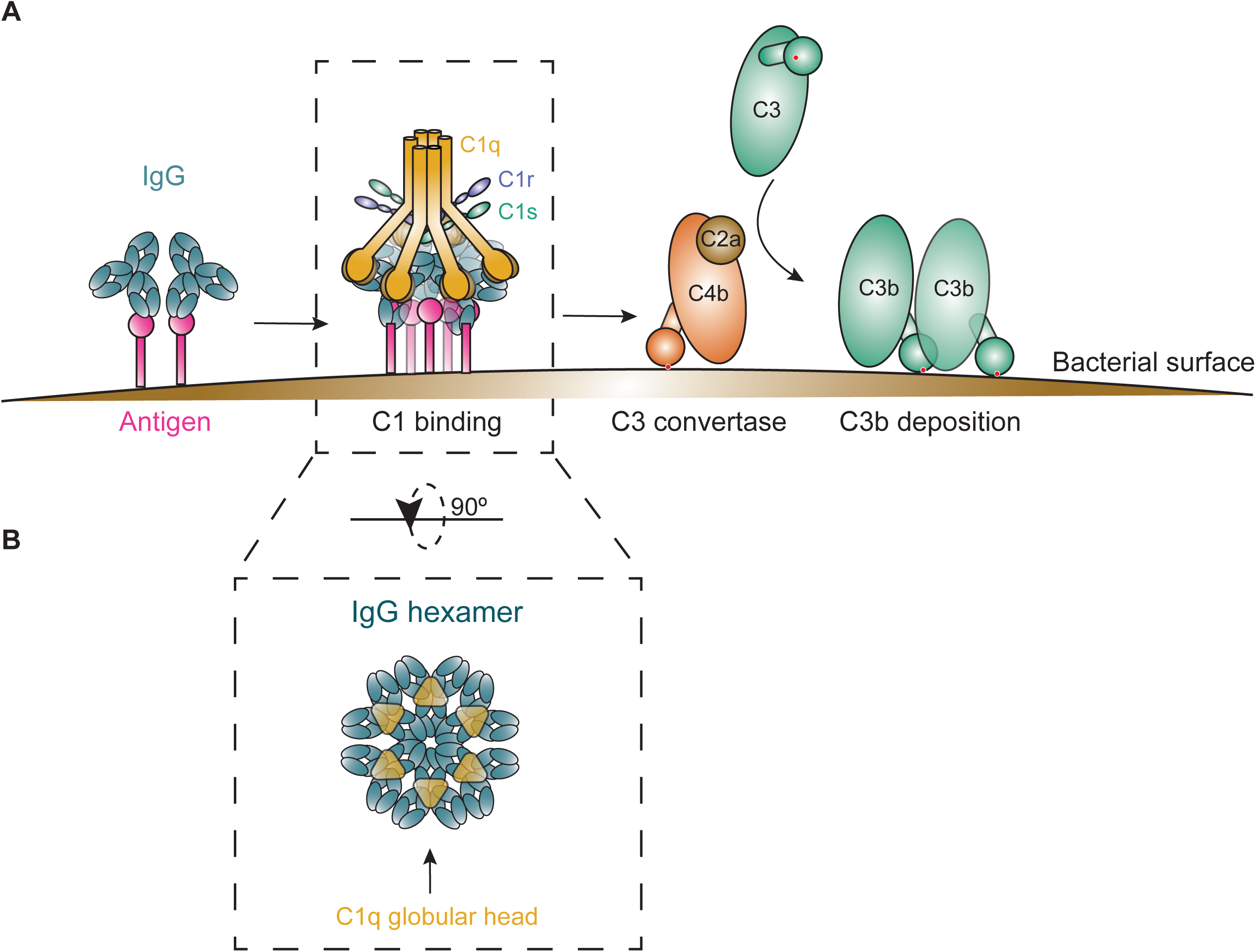
Graphical representation of complement classical pathway activation. A Overview of complement classical pathway initiation. The fully assembled C1 complex binds to immune complexes on the bacterial surface, subsequently, its attached C1r/C1s proteases are activated and cleave C4 and C2 to generate C4b2a (C3 convertase). The C3 convertase cleaves C3 into C3b, which displays a previously hidden thioester that covalently binds to the bacterial surface. B The IgG hexameric rings, or IgG hexamers, are held together by non-covalent Fc-Fc interactions and form the perfect docking platform for complement C1.

**Figure EV2.**
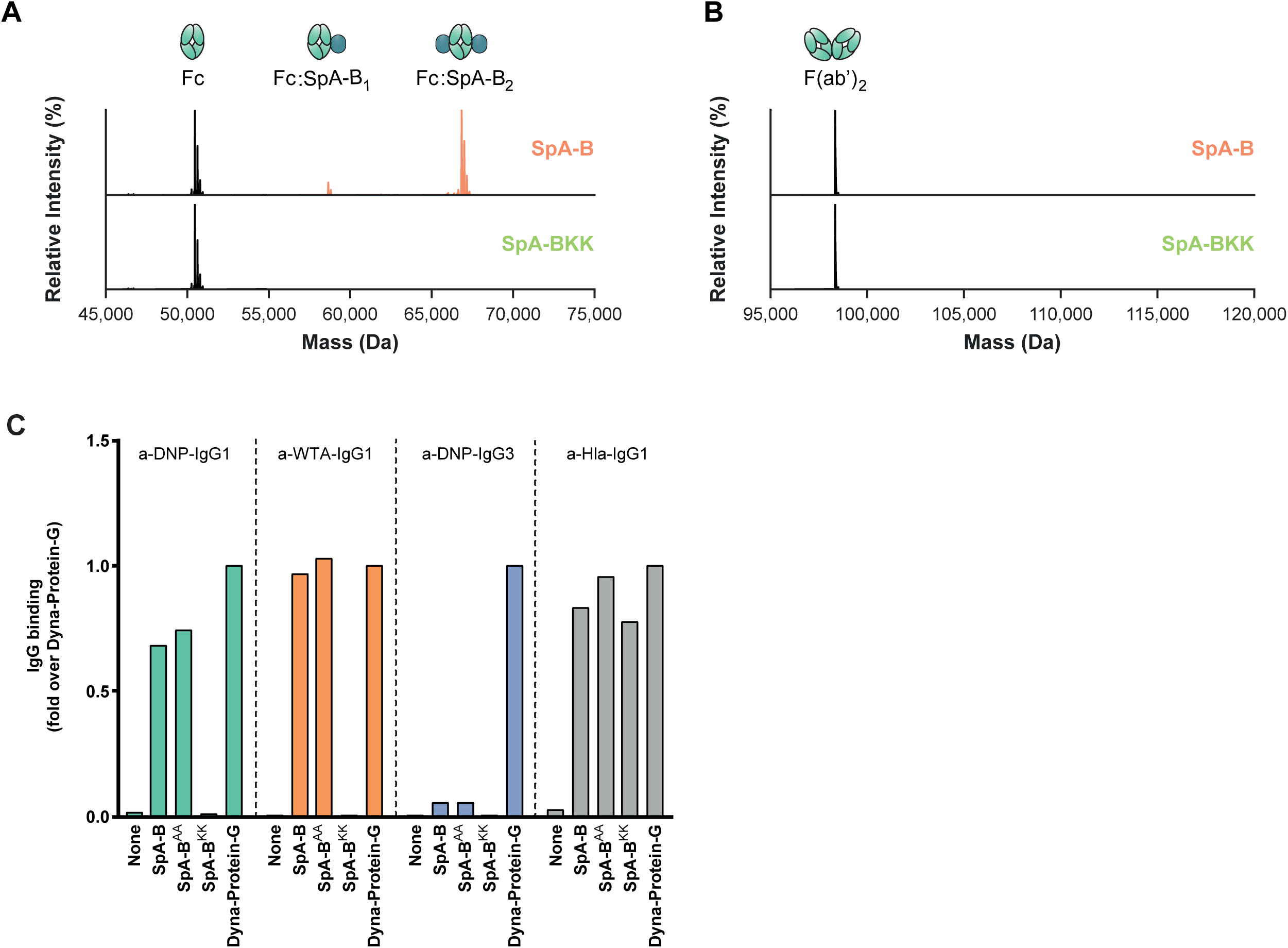
Human anti-DNP and anti-WTA antibodies only bind to SpA via their Fc region. A, B Deconvoluted native mass spectra of the Fc (A) and F(ab’)_2_ (B) molecules of anti-CD52 IgG1, resulting from IdeS digestion, in absence (black) or presence of SpA-B (orange) or SpA-B^KK^ (green). SpA-B binds to the Fc region, but not to Fab region. SpA-B^KK^ does not bind to the Fc or Fab region. C Binding of anti-DNP IgG1, anti-DNP IgG3, anti-WTA IgG1 and anti-Hla IgG1 to beads coated with His-tagged recombinant SpA-B wild-type (wt), SpA-B^AA^ (that only binds to IgG-Fc region), SpA-B^KK^ (that only binds to IgG-Fab region) or to Protein-G beads. Antibody binding was detected by the use of directly labeled IgG or Alexa Fluor^647^-conjugated goat F(ab’)_2_ anti-human kappa and was measured by flow cytometry. The anti-Hla IgG1 (VH3 family antibody) served as a control for Fab binding to SpA-B^KK^. Protein-G beads served as universal IgG1 and IgG3 binding control.

**Figure EV3.**
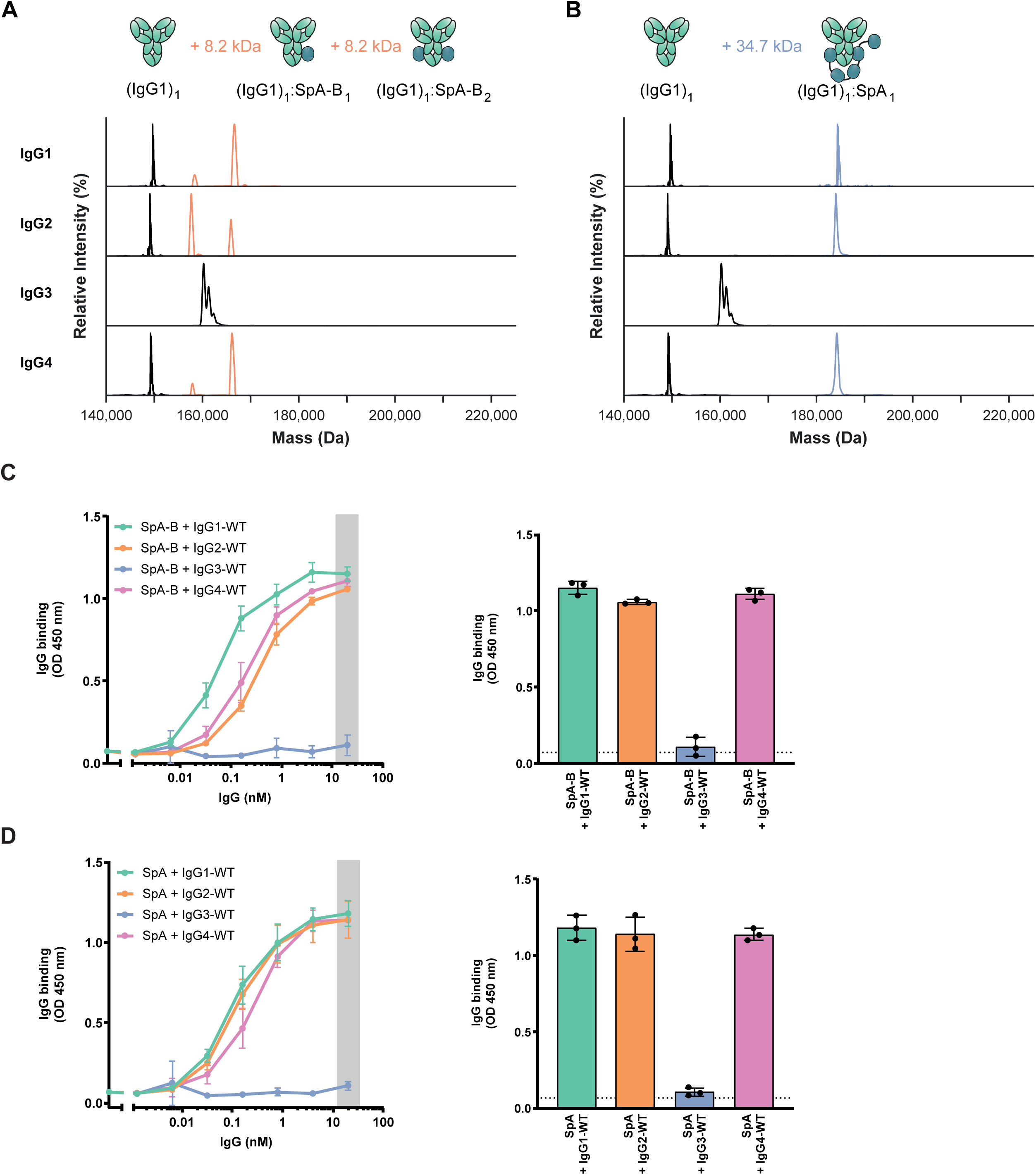
Human IgG1, IgG2 and IgG4 bind to SpA, but not IgG3. A, B Deconvoluted native mass spectra revealing the binding stoichiometry of SpA-B and SpA with IgG. Incubation with SpA-B (orange) results in mass shifts corresponding to SpA-B binding to IgG1, IgG2 and IgG4 with up to 2:1 stoichiometry (A) whereas SpA (blue) binds principally with 1:1 stoichiometry to the same IgG subclasses (B). Satellite peaks observed in the mass spectra of IgG3 result from the presence of additional O-glycosylation sites on the IgG3 hinge region. C, D Binding of the full concentration range of human IgG subclasses to SpA-B (C) or SpA (D) coated on a 96-well plate. IgG binding was detected with HRP-labeled goat F(ab’)_2_ anti-human kappa. Bars represent the same data for the 20 nM IgG concentration only and the black dotted line shows the background absorbance from wells that were not incubated with IgG.

**Figure EV4.**
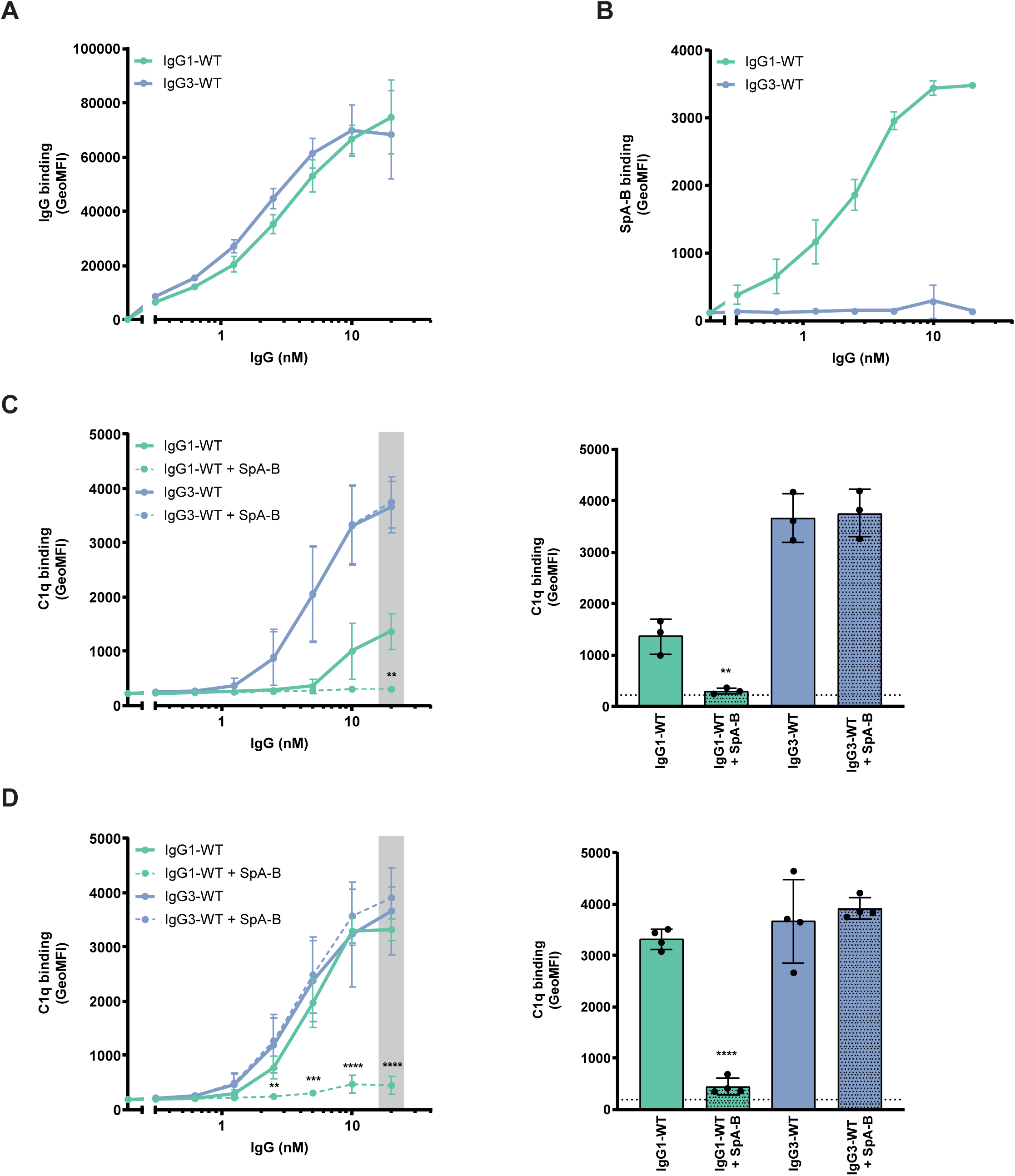
The binding of SpA-B to antigen-bound IgG decreases C1q binding on target surfaces even if C1q is incubated before SpA-B. A Binding of anti-DNP IgG1-WT and IgG3-WT to DNP-coated beads, detected with Alexa Fluor647-conjugated goat F(ab’)_2_ anti-human kappa by flow-cytometry. B Binding of SpA-B to anti-DNP IgG1-WT and IgG3-WT bound to DNP-coated beads, detected with anti-HexaHistidine chicken antibody by flow cytometry. C, D C1q binding on IgG1-WT and IgG3-WT bound to DNP-coated beads, detected with FITC-conjugated rabbit F(ab’)_2_ anti-human C1q by flow-cytometry. Buffer (solid lines) or SpA-B (dotted lines) was added to the beads only after C1q (C) or C1 complex (D). Bars represent the same data for the 20 nM IgG concentration only and the black dotted line shows the background fluorescence from beads that were not incubated with IgG. Data are presented as geometric means ± SD of three (C) or four (D) independent experiments. Statistical analysis was performed using an unpaired two-tailed t-test to compare buffer and SpA-B conditions and showed when significant as **P ≤ 0.01, ***P≤ 0.001 and ****P ≤ 0.0001.

**Figure EV5.**
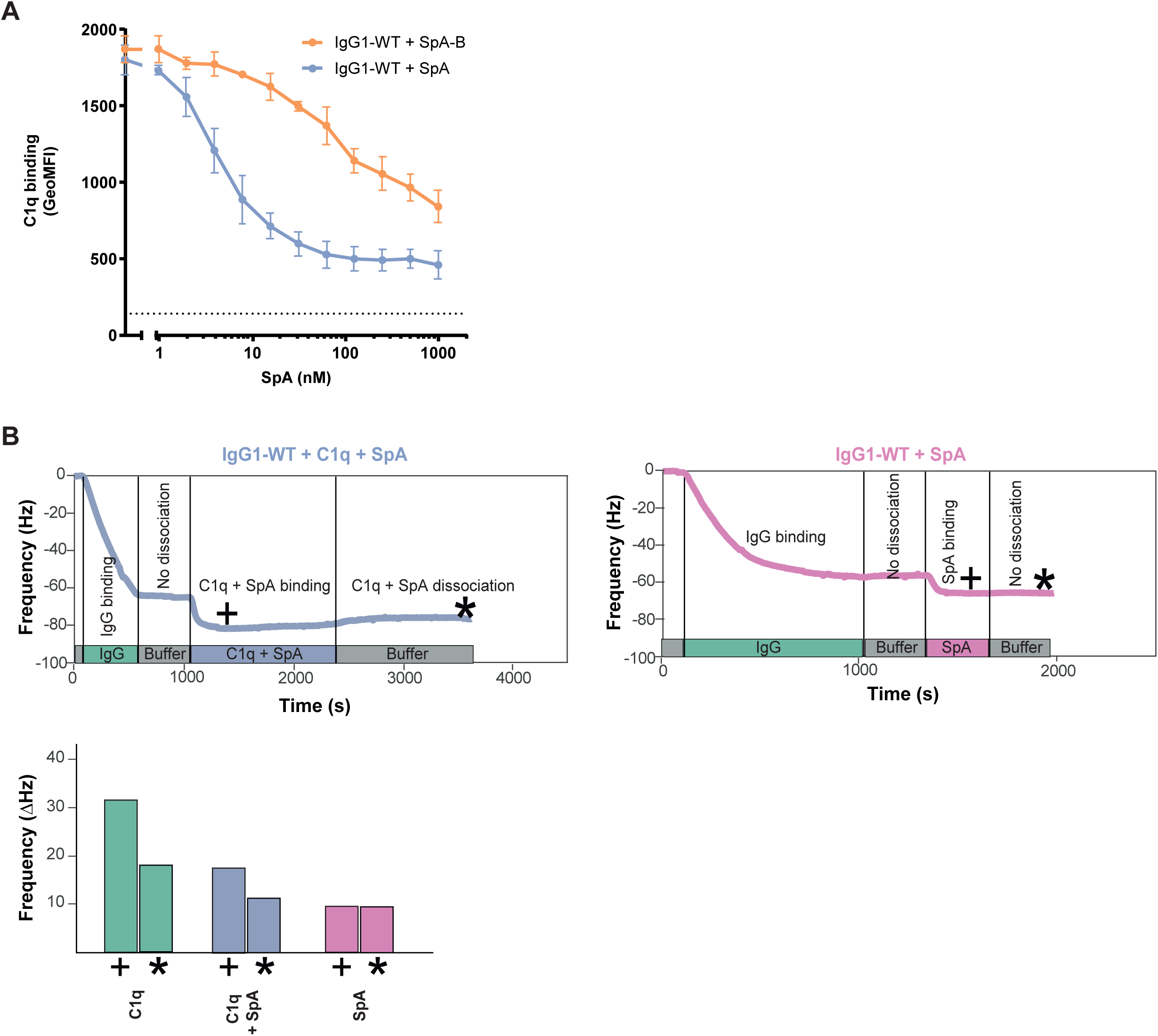
SpA inhibits binding of C1q to antigen-bound IgGs on target surfaces. A C1q binding on anti-DNP IgG1-WT bound to DNP-coated beads after incubation of C1 complex with a concentration range of SpA-B (orange) or SpA (blue), detected with FITC-conjugated rabbit F(ab’)_2_ anti-human C1q antibody by flow cytometry. The black dotted line shows the background fluorescence from beads that were not incubated with IgG. Data are presented as geometric means ± SD of at least three independent experiments. B, C QCM sensorgram of C1q binding to anti-DNP IgG1-WT bound to DNP-decorated lipid bilayers in the presence of SpA (B) and SpA binding (in the absence of C1q) to anti-DNP IgG1-WT bound to DNP-decorated lipid bilayers (C). Bars represent the respective equilibrium level (+) and the level at the end of the dissociation phase (*) (C1q sensorgram is presented in Fig 5A).

**Figure EV6.**
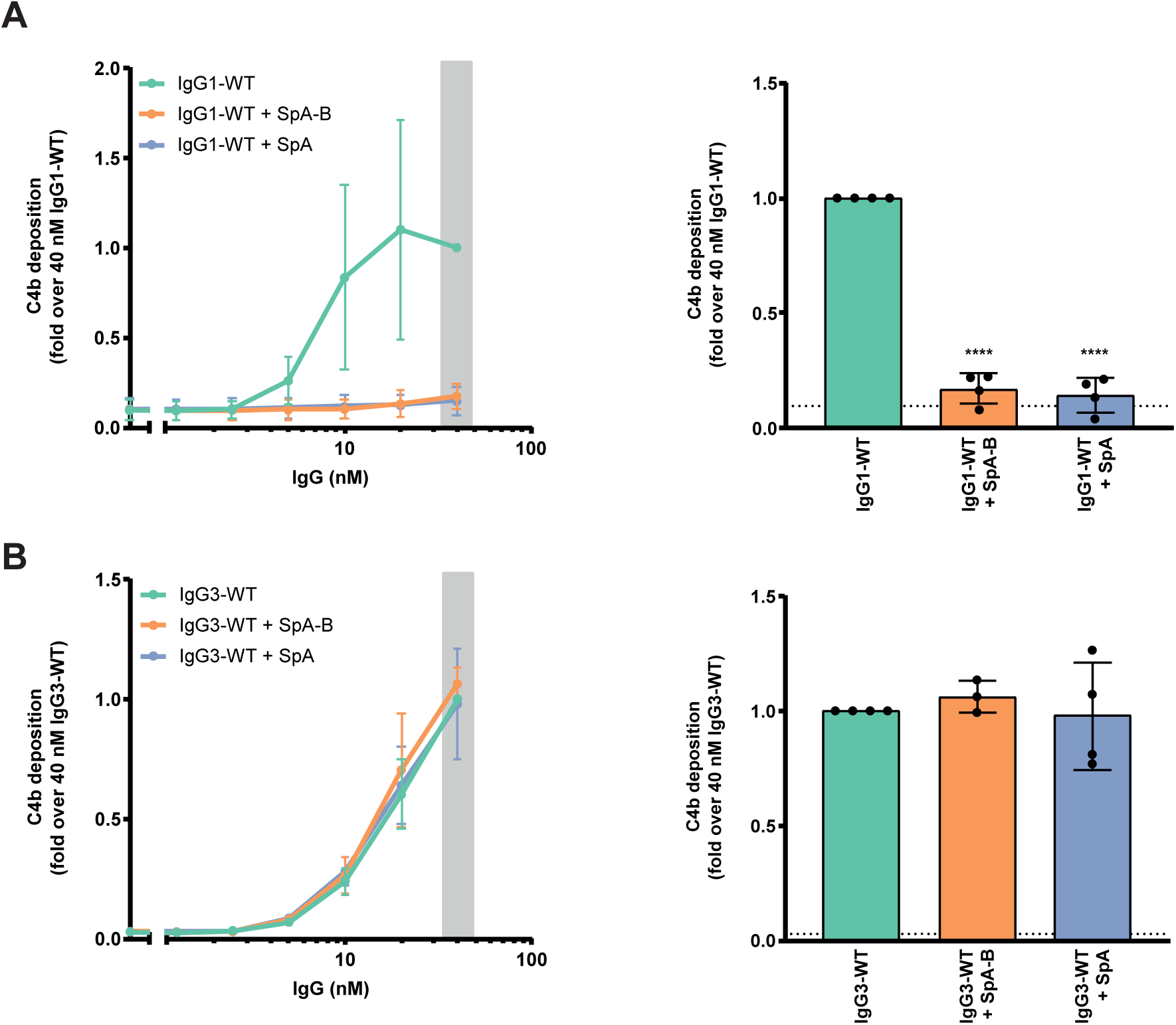
SpA decreases IgG-mediated C4b deposition on *S. aureus*. A, B C4b deposition on NewmanΔ*spa*/*sbi* surface after bacteria incubation with IgG1-WT (A) or IgG3-WT (B), 1% ΔIgG/IgM human serum and buffer (green), SpA-B (orange) or SpA (blue), detected with a monoclonal murine anti-human C4d antibody by flow cytometry. Bars represent the same data for the 40 nM IgG concentration only and the black dotted line shows the background fluorescence from bacteria that were not incubated with IgG. Data are presented as fold change over the 40 nM IgG concentration control ± SD of at least three independent experiments. Statistical analysis was performed using one-way ANOVA to compare buffer condition with SpA-B and SpA conditions and showed when significant as ****P ≤ 0.0001.

**Table EV1.** Theoretical and experimental molecular weight values of proteins analyzed by native MS. The mass deviations of ≈ 2.5 kDa for the IgGs are due to the N-glycosylations in the Fc region, whereas the 42 kDa mass difference for C1q is expected due to extensive glycosylation of the □, □ and □ subunits.

| Species | MW<sub>theoretical</sub> (Da) | MW<sub>experimental</sub> (Da) | ΔMW (Da) |
| --- | --- | --- | --- |
| SpA-B | 8,320 | 8,188 | -132* |
| SpA-BKK | 8,320 | 8,188 | -132* |
| SpA | 34,660 | 34,663 | 3 |
| IgG1 | 146,995 | 149,217 | 2,222 |
| IgG1-RGY | 147,057 | 149,831 | 2,774 |
| IgG2 | 146,428 | 149,065 | 2,636 |
| IgG2-RGY | 146,490 | 149,137 | 2,647 |
| IgG3 | 157,219 | 160,113 | 2,894 |
| IgG3-RGY | 157,281 | 159,929 | 2,647 |
| IgG4 | 146,544 | 149,181 | 2,637 |
| IgG4-RGY | 146,606 | 149,237 | 2,631 |
| C1q | 421,437 | 463,591 | 42,153 |

\* Corresponding to the deletion of Met1 in the protein sequence.

**Table EV2.**
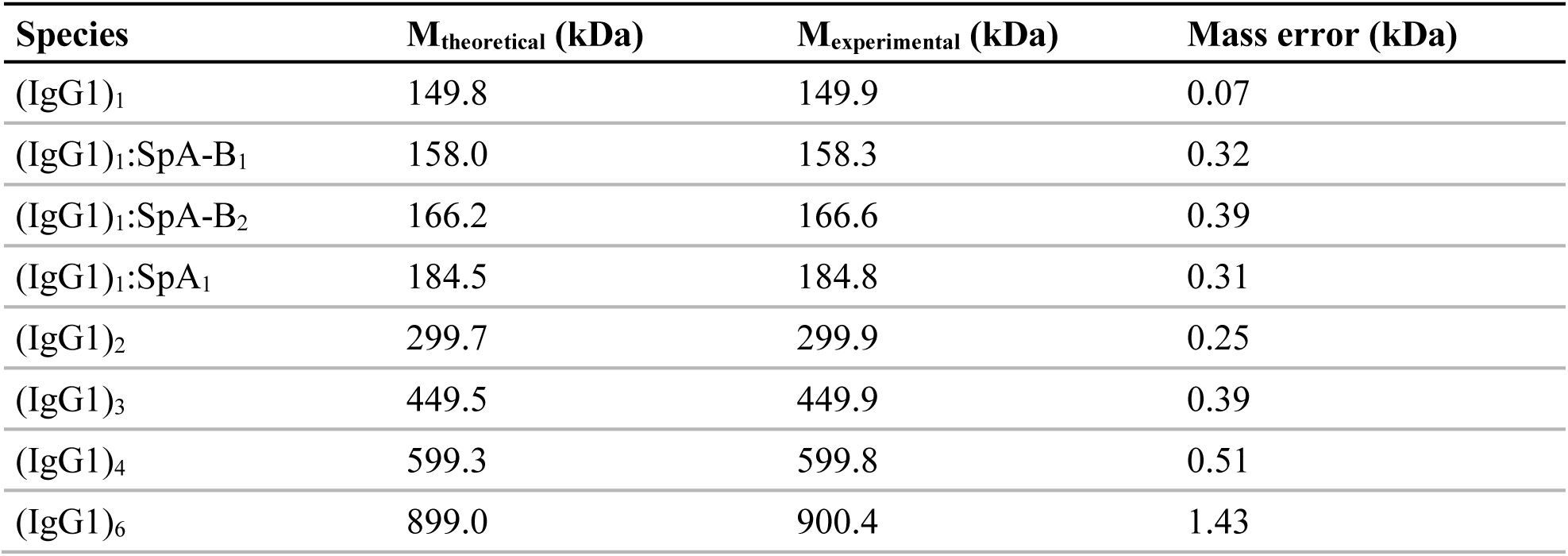
Theoretical and experimental mass values of proteins complexes related to Fig 2A, native mass spectra showing the effect of SpA-B and SpA on IgG1-RGY hexamerization. The theoretical mass of the complex, as calculated using the measured masses of individual subunits, is compared to the experimental mass. For these experiments, instrumental parameters were optimized for the quantification of (IgG)_6_ complexes.

**Table EV3.**
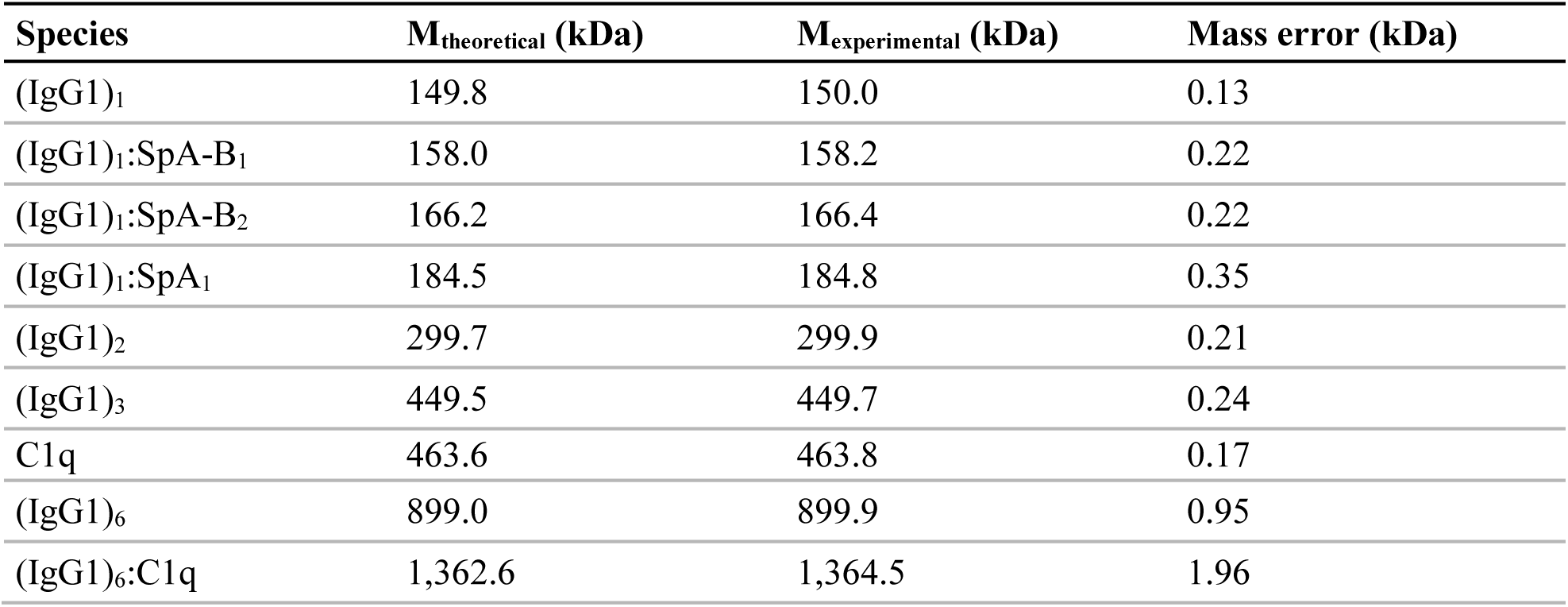
Theoretical and experimental mass values of proteins complexes related to Fig 3A, native mass spectra showing the effect of SpA-B and SpA on (IgG1-RGY)_6_:C1q assembly. The theoretical mass of the complex, as calculated using the measured masses of individual subunits, is compared to the experimental mass. For these experiments, instrumental parameters were optimized for the quantification of (IgG)_6_:C1q complexes.

| **Species** | **M<sub>theoretical</sub> (kDa)** | **M<sub>experimental</sub> (kDa)** | **Mass error (kDa)** |
| --- | --- | --- | --- |
| (IgG1)<sub>1</sub> | 149.8 | 150.0 | 0.13 |
| (IgG1)<sub>1</sub>:SpA-B<sub>1</sub> | 158.0 | 158.2 | 0.22 |
| (IgG1)<sub>1</sub>:SpA-B<sub>2</sub> | 166.2 | 166.4 | 0.22 |
| (IgG1)<sub>1</sub>:SpA<sub>1</sub> | 184.5 | 184.8 | 0.35 |
| (IgG1)<sub>2</sub> | 299.7 | 299.9 | 0.21 |
| (IgG1)<sub>3</sub> | 449.5 | 449.7 | 0.24 |
| C1q | 463.6 | 463.8 | 0.17 |
| (IgG1)<sub>6</sub> | 899.0 | 899.9 | 0.95 |
| (IgG1)<sub>6</sub>:C1q | 1,362.6 | 1,364.5 | 1.96 |

**Table EV4.**
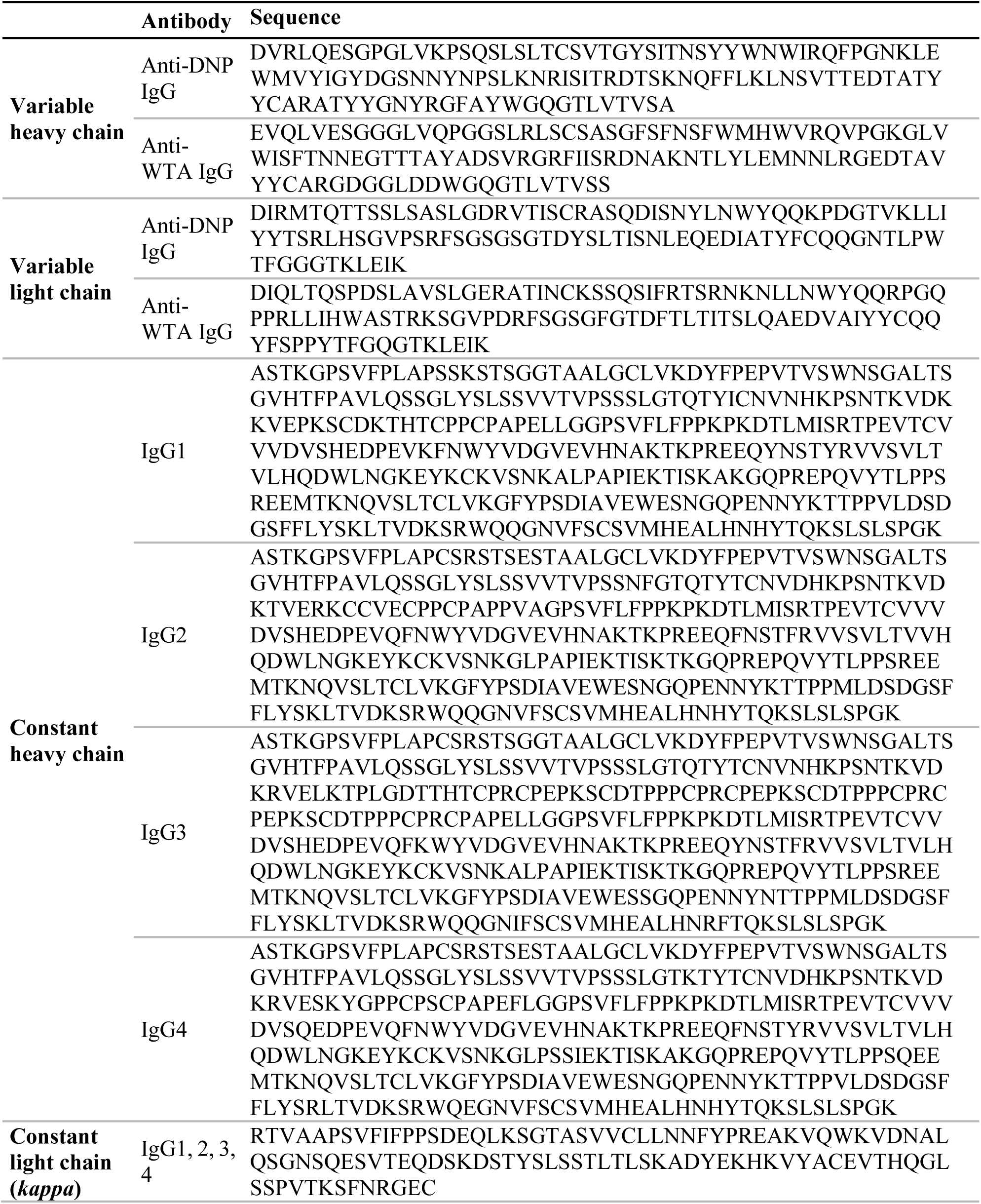
Amino acid sequence of the variable and constant heavy and light chains of antibodies produced in this study (Gonzalez *et al*, 2003; Lehar *et al*, 2015).

**Table EV5.**
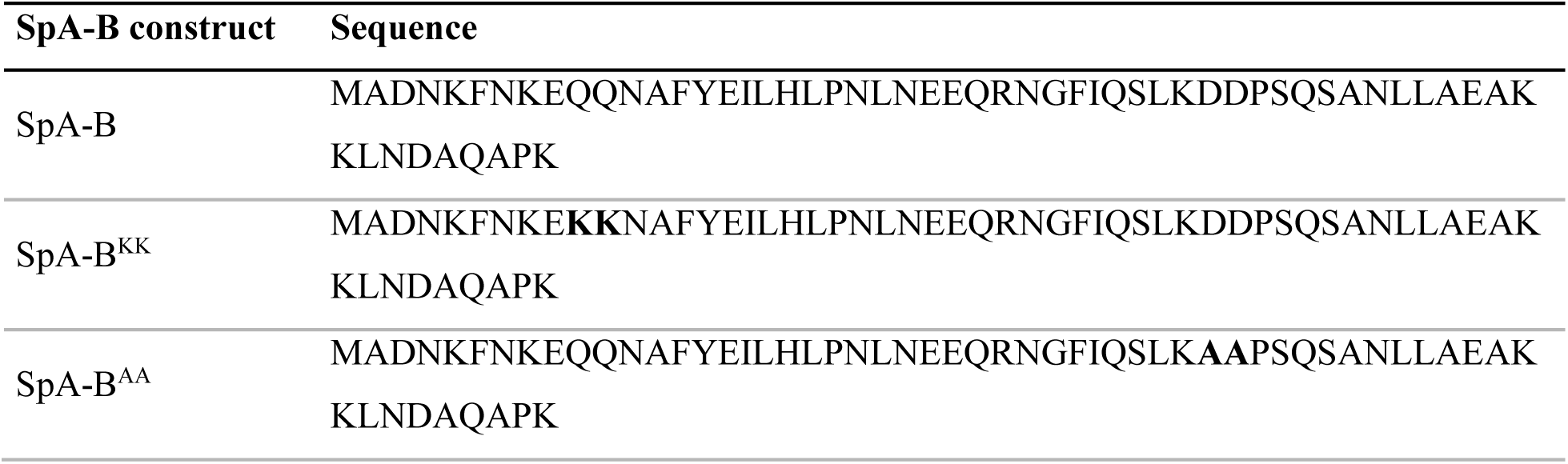
Amino acid sequence of SpA-B constructs. The amino acid substitutions that are highlighted in bold abrogate binding of SpA-B^KK^ and SpA-B^AA^ to Fc and Fab domain of human IgG, respectively.

